# ULK4 and Fused/STK36 interact to mediate assembly of a motile flagellum

**DOI:** 10.1101/2022.03.06.483169

**Authors:** Ciaran J. McCoy, Humbeline Paupelin-Vaucelle, Peter Gorilak, Tom Beneke, Vladimir Varga, Eva Gluenz

## Abstract

Unc-51-like kinase (ULK) family serine-threonine protein kinase homologs have been linked to the function of motile cilia in diverse species. Mutations in Fused/STK36 and ULK4 in mice resulted in hydrocephalus and other phenotypes consistent with ciliary defects. How either protein contributes to the assembly and function of motile cilia is not well understood. Here we studied the phenotypes of *ULK4* and *Fused* gene knockout (KO) mutants in the flagellated protist *Leishmania mexicana*. Both KO mutants exhibited a variety of structural defects of the flagellum cytoskeleton. Biochemical approaches indicate spatial proximity of these proteins and indicates a direct interaction between the N-terminus of LmxULK4 and LmxFused. Both proteins display a dispersed localisation throughout the cell body and flagellum, with enrichment near the flagellar base and tip. Fused/STK36 was previously shown to localise to mammalian motile cilia and we demonstrate here that ULK4 also localises to the motile cilia in mouse ependymal cells. Taken together these data suggest a model where the pseudokinase ULK4 is a positive regulator of the kinase Fused/STK36 in a pathway required for stable assembly of motile cilia.

**Summary Statement:** Knockout phenotypes in *Leishmania*, and confirmation of ULK4 ciliary localisation in mouse, show ULK4 and Fused/STK36 interact in a conserved pathway for stable assembly of motile cilia.

## Introduction

Serine-threonine protein kinases of the Unc-51-like kinase (ULK) family are found across eukaryotic species, with the best-characterised members found in metazoans and plants. In mammals the family has four active kinases (ULK1, ULK2, ULK3 and Fused/STK36) and one pseudokinase ULK4. The active kinases function in multiple well-studied biological processes: ULK1 and ULK2 act in autophagy pathways in starvations responses (Lee, 2011), ULK3 is a regulator of the sonic hedgehog signalling pathway (Maloverjan et al., 2010) and Fused kinase was first identified in *Drosophila* (Preat et al., 1990) as a kinase of Hedgehog (Hh) signalling pathway. Unexpectedly, studies on the mouse Fused homolog, also named Serine/threonine-protein kinase 36 (STK36), showed that *fused* knockout mice did not show any defects in Hh signalling (Chen et al., 2005; Merchant et al., 2005). Instead, it was found that these mice developed hydrocephalus and respiratory infections and died a few weeks after birth (Merchant et al., 2005). This phenotype, together with the finding that Fused was highly expressed in ependymal cells and nasal epithelium, pointed to a function in cells with motile cilia.

Cilia and flagella are cellular appendages formed by a microtubule axoneme, which originates at the basal body and has more or less extensive structural elaborations, depending on its specific function within a cell type. A key distinction is between motile cilia/flagella, which typically have a so-called 9+2 axoneme composed of a pair of singlet microtubules (central pair; CP) surrounded by a ring of nine doublet microtubules. The latter are decorated with axonemal dyneins and other protein complexes such as radial spokes (RSP) required for bending. They provide one of the main modes of eukaryotic cell motility, as used by many protists, unicellular plants and gametes of diverse species including mammalian sperm cells. Cells extending motile cilia are also found within tissues where they generate directional fluid flow required for development and normal physiology (Ringers et al., 2020), e.g. ciliated ependymal cells of the brain ventricles and spinal canal help the circulation of cerebrospinal fluid. Primary cilia by contrast have a simpler axoneme structure consisting of 9 doublet microtubules and they transduce chemical and mechanical signals from the environment (Goetz and Anderson, 2010). Cilia and flagella construction relies on a conserved intraflagellar transport (IFT) system that shuttles cargo, including axonemal constituents, from the basal body to the axoneme tip and back (Lechtreck, 2015).

Wilson et al. (2009) Investigated whether mammalian Fused was required for the Hh responses mediated through primary cilia and found that this was not the case: *Fu*^*-/-*^ mice were capable of constructing normal primary cilia. By contrast, *Fu*^*-/-*^ mice had motile cilia defects with 60% showing an abnormal ultrastructure (40% of cilia had no central pair) and misalignment, which resulted in uncoordinated ciliary beating. Unlike the situation in mice, morpholino-mediated knockdown of *Fu* in Zebrafish did affect Hh signalling, and in addition, the morphants also had structurally and functionally disrupted cilia in Kupffer’s vesicle, which are required in development to generate a directional fluid flow to establish the body’s left-right axis. The effect on the cilia was independent of Hh signalling, indicating that this kinase acts in multiple conserved pathways in different lineages (Wilson et al., 2009). Studies on oviduct cilia provided further evidence for an important role of mammalian Fused/STK36 in motile cilia. The majority of oviduct cilia of *Fu*^*-/-*^ mice had an abnormal axoneme structure (Nozawa et al., 2013): cilia were misaligned and a majority of these oviduct cilia lacked central pair microtubules, within tissues that also produced some cilia with a normal 9+2 architecture. Mouse sperm flagella function were also affected by conditional deletion of *Fu*, showing motility defects associated with infertility (Nozawa et al., 2014). In the sperm cells the 9+2 microtubule structure of the axoneme was maintained but they exhibited a defect in the structure of the manchette, a transient microtubule structure required for correct shaping of the sperm head.

In humans, primary ciliary dyskinesia (PCD) is a disease caused by defects in motile cilia, which can result from varied genetic causes. Fused/STK36 loss-of-function mutations were found in a PCD patient (Edelbusch et al., 2017) where most motile cilia were normal but ~5% had abnormal microtubule numbers and cilia mis-orientation affected their movement. The Fu protein itself was localised along the length of the motile cilium (Edelbusch et al., 2017; Nozawa et al., 2013), consistent with the genetic evidence for a role of Fused/STK36 in motile cilia. One study suggested it was sited between the radial spoke heads and the central pair microtubules (Edelbusch et al., 2017). Biochemical evidence for an interaction between mouse Fu and the central pair-associated protein Spag16/PF20 and several other ciliary proteins led to a model where Fu and the kinesin-like protein Kif27 interact to promote CP assembly in motile cilia of mammals (Wilson et al., 2009).

Less functional information is available for the pseudokinase UKL4, but, like for Fused/STK36, there is converging evidence for a function associated with motile cilia. In a mouse knockout and phenotyping screen, ULK4 was identified in a cohort of genes for which disruption caused hydrocephalus (Vogel et al., 2012). This gene cohort also included Fused/STK36 and Kif27. In *Ulk4* hypomorph mutant mice, in which *Ulk4* expression was reduced, ependymal cells had abnormal axonemes and disorganised cilia (Liu et al., 2016). Abnormal structures included 9+0 and other deviations from normal 9+2 microtubule numbers, while some cilia had normal structures. The possibility was raised that UKL4 regulates different processes of ciliogenesis. RNA-seq showed alterations in cilia related transcripts and indicated that UKL4 regulated Foxj1 transcription factor pathways involved in ciliogenesis.

The documented links between ULK4 disruption and human diseases prompted the elucidation of the ULK4-ATPγS complex structure and mapping of the molecular ‘environment’ by proximity labelling (Preuss et al., 2020). The proximity interaction network uncovered five kinases, including Fused/STK36, and proteins associated with a range of different functions, amongst them ciliary and microtubule-associated proteins.

Orthologs of Fused/STK36 and UKL4 are present in the kinetoplastids, a group of flagellated protozoans that are well-studied on grounds of their medical importance as causative agents of neglected parasitic diseases. In *Trypanosoma brucei*, FCP6/TbFused and FCP5/TbULK4 localised to the flagella connector (Varga et al., 2017), a mobile transmembrane structure at the tip of the newly growing flagellum that connects it to the existing flagellum during a defined phase in the growth cycle (Moreira-Leite et al., 2001). Deletion of *LmxFused* and *LmxULK4* in *Leishmania mexicana*, in a ‘kinome’-wide knockout (KO) screen yielded cells with shorter flagella and reduced motility (Baker et al., 2021). We independently found *LmxUKL4* and *LmxFused* in a screen of flagellar protein KO mutants with reduced motility. Here, we conducted a detailed analysis of these null mutant phenotypes and found that *LmxULK4* and *LmxFused* KOs both present with the same spectrum of morphological defects, affecting length and ultrastructure of the flagellar axoneme. This indicates a conserved role for Fused/STK36 or ULK4 in ciliogenesis and cilia function. Our data support a model whereby these two proteins directly interact with each other to promote stable assembly of motile flagella in evolutionarily distant lineages. Consistent with this model, we also show that the mammalian ULK4 protein localises to the motile cilium of mouse ependymal cells.

## Results

### LmxULK4 and LmxFused localise to both the Leishmania cell body and motile flagellum

*LmxULK4* and *LmxFused* encode UNC-51-like proteins that are orthologs of animal ULK4 and Fused/STK36, respectively (Fig 1A). Like its mammalian orthologs, LmxULK4 represents a catalytically inactive pseudokinase, as supported by the lack of the catalytically important Lysine residue within the typically highly conserved ‘VAIK’ motif (see Fig 1B). We assessed protein localisation by inserting an mNeonGreen (mNG) fusion tag at the endogenous gene locus (tagging both alleles of the gene) and imaging the fusion proteins in live cells. Both proteins localised to the *Leishmania* promastigote cell body and motile flagellum throughout the cell cycle (Fig 1C-E). Individual cells (both dividing and non-dividing) often displayed enriched signal at the flagellum tip, base or both but no evidence was found for a dominant signal uniquely localised at the tip of the newly growing flagellum, where the orthologous *T. brucei* proteins are localised (Varga et al., 2017). The LmxFused::mNG and LmxULK4::mNG fluorescent signal was absent in cells extracted with a nonionic detergent (S1 Fig), suggesting that neither protein is tightly attached to the insoluble cytoskeleton.

**Fig 1.**
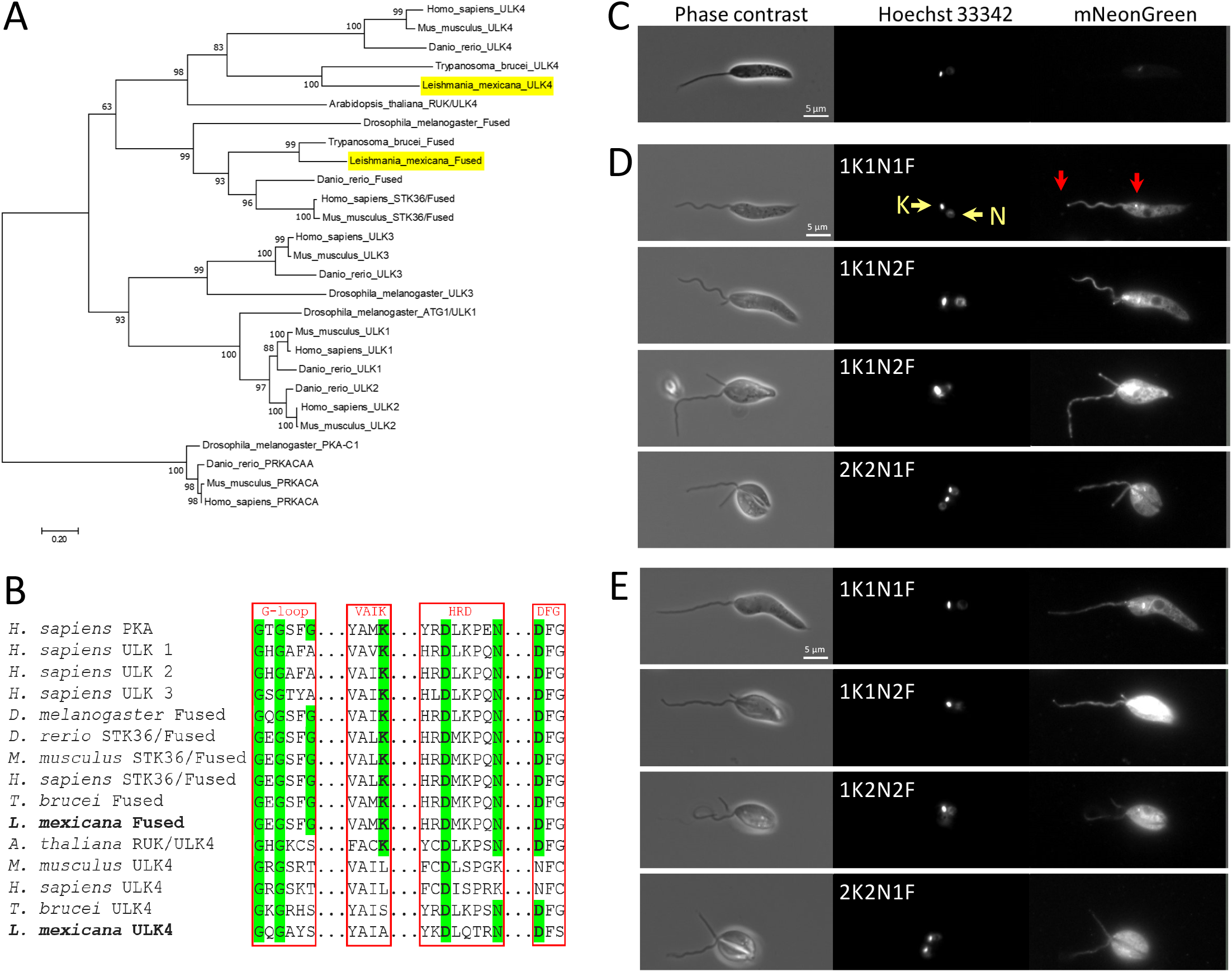
UNC-51 like kinase orthologs in L. mexicana and their subcellular localisation. **(A)** Position of *L. mexicana* (Lmx) Fused and *Lmx*ULK4 in relation to UNC-51 like kinases. Maximum likelihood tree based on the LG (+*G*+*I*) model of evolution and with 500 bootstrap replicates (MEGA 7 (Kumar et al., 2016); sequences used: *H. sapiens* ULK1 (O75385), ULK2 (Q8IYT8), ULK3 (Q6PHR2), ULK4 (Q96C45), STK36/Fused (Q9NRP7), PRKACA (P17612); *M. musculus* ULK1 (O70405), ULK2 (Q9QY01), ULK3 (Q3U3Q1), ULK4 (Q3V129), STK36/Fused (Q69ZM6), PRKACA (P05132); *D. rerio* ULK1 (F1R9T2), ULK2 (X1WEA3), ULK3 (A4IG43), ULK4 (A0A0R4IA69), Fused (A8WFS2), PRKACAA (A3KMS9); *D. melanogaster* ATG1/ULK1 (Q9VU14), ULK3 (Q9VHF6), Fused (P23647), PKA-C1 (P12370); *A. thaliana* RUK/ULK4 (F4JY37), with UniProt IDs in brackets (UniProt, 2021). *T. brucei* FCP5/ULK4 (Tb927.11.8150), FCP6/Fused (Tb927.11.4470); *L. mexicana* ULK4 (LmxM.28.0620), Fused (LmxM.13.0440) with TritrypDB GeneIDs (Amos et al., 2021) in brackets. **(B)** Kinase motif (‘G-loop’, ‘VAIK’, ‘HRD’ and ‘DFG’ motifs) focused multiple sequence alignment. Catalytically relevant amino acid residues are highlighted in green, with the residues that form the canonical catalytic triad also highlighted in bold. **(C-E)** Fluorescence micrographs of *L. mexicana* promastigote forms, expressing either no fusion protein (parental control, C), LmxFused::mNG (D) or LmxULK4::mNG (E). The micrographs show different cell cycle stages, as assessed by the number of kinetoplasts (K), nuclei (N), and flagella (F). Nuclear and kinetoplast DNA were labelled with Hoechst 33342. Red arrows point to the base and tip of the flagellum.

### Loss of LmxULK4 or LmxFused leads to failure of normal motile flagellum assembly

To analyse loss-of-function phenotypes, CRISPR/Cas9 was used to knock out (KO) *LmxULK4* and *LmxFused*, respectively (S2 Fig). Measurement of swimming speed showed that this was significantly reduced in both KO lines (Fig 2A), relative to parental controls, consistent with a lack of a long flagellum in the majority (~90-95%) of cells in these KO populations (n≥500; Fig 2B, D). The visible flagella that were still present in a minority of the KO cells were shorter than those of the parental controls (n≥27; Fig 2C; one-way ANOVA with the Šídák correction for multiple comparisons). Moreover, although these were not completely paralysed, they did not enable cell propulsion but rather facilitated a small proportion of cells to twitch at the bottom of their culture dish when observed *in vitro*. Despite these striking changes in morphology, the KO mutants only exhibited a very slight increase in doubling time (S3 Fig). Importantly, the flagellum-associated phenotypes reported here were completely rescued via re-introduction of the target gene on an episomal addback plasmid for the Δ*LmxFused* cell line and partially rescued for Δ*LmxULK4* (Fig 2A-C). A dual KO of both genes (Δ*LmxULK4* / Δ*LmxFused*) resulted in flagellar phenotypes that were indistinguishable to either individual Δ*LmxULK4* and Δ*LmxFused* cell line (Fig 2A-D).

**Fig 2.**
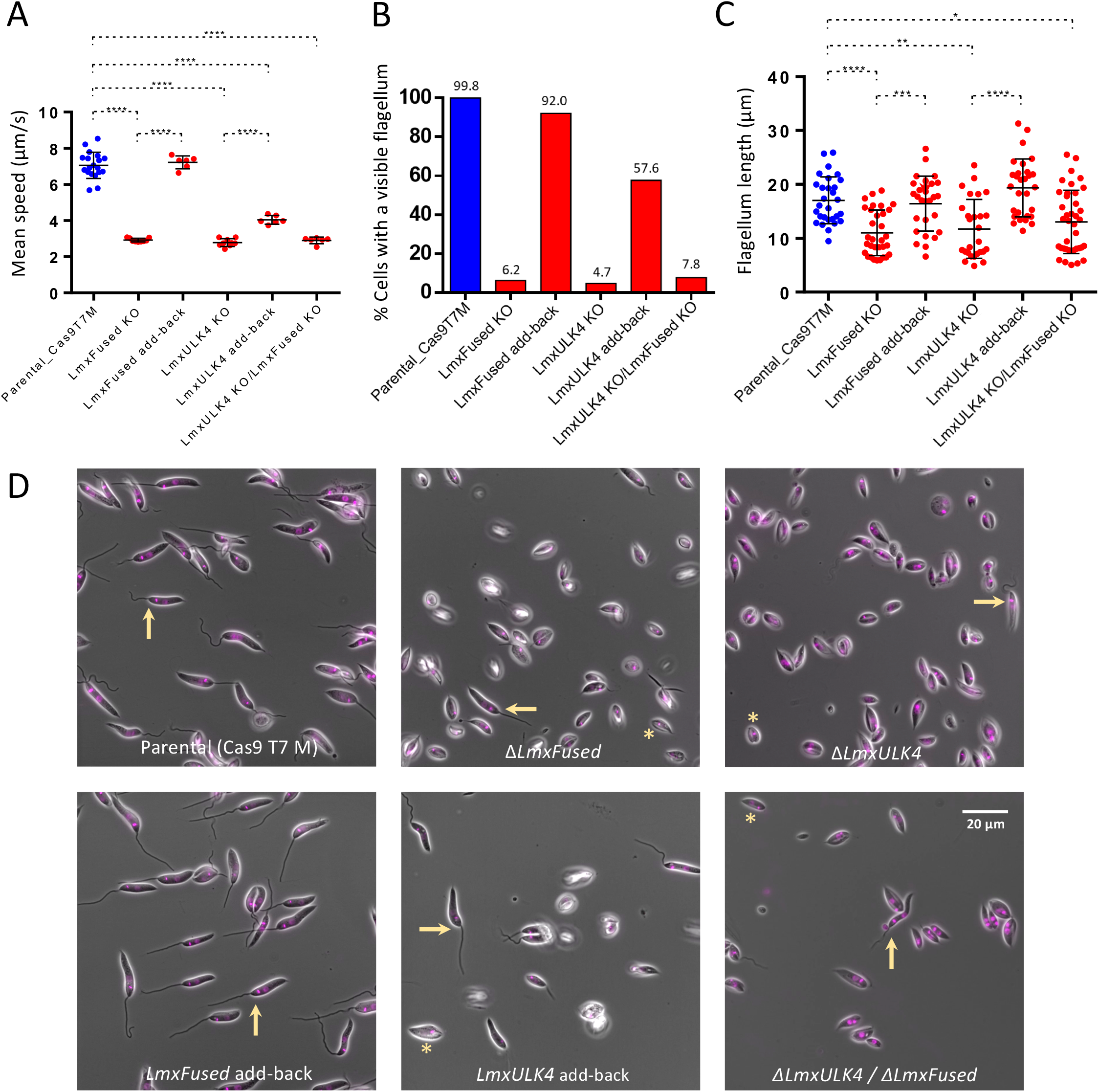
Deletion of LmxULK4 and LmxFused results in loss of motile flagella. **(A)** Swimming speed of the parental *L. mexicana* promastigote cells, *LmxULK4* and *LmxFused* KO mutants and respective add-back cell lines. Each dot represents the average speed of promastigotes from one motility assay. **(B)** Proportion of cells with a visible external flagellum. **(C)** Measurement of flagellar length, for cells that had an external flagellum. Each dot represents an individual cell. **(D)** Morphology of the parental cell line, *LmxULK4* and *LmxFused* KOs and episomal add-back cell lines. Images are composites of the phase contrast channel and a fluorescence channel showing nuclear and kinetoplast DNA labelled with Hoechst 33342. Arrows point to examples of cells with a visible external flagellum. An asterisk highlights example cells that lack a visible external flagellum.

### ΔLmxULK4 and ΔLmxFused mutants display a range of structural defects in the flagellum

These data are consistent with a role of *LmxFused* and *LmxULK4* in the assembly of a motile flagellum (Baker et al., 2021). A striking feature of the KO phenotypes was that within clonal populations of apparently aflagellate cells, a small but stable proportion of cells did still possess a long flagellum. This suggests that LmxULK4 or LmxFused are not strictly essential for the formation of a flagellum, but their absence prevented the reliable assembly of functional motile flagella in most cells. To understand which step of flagellum biogenesis failed in these mutants, the structure of the flagella in the Δ*LmxFused* and Δ*LmxULK4* populations was examined in detail. The axonemal radial spoke protein 11 (RSP11; LmxM.09.1530) and the central pair associated protein PF16 (LmxM.20.1400) were chosen as reporter proteins for key structures of the motile axoneme and a cell line was generated simultaneously expressing RSP11::eYFP and PF16::mStrawberry (Pf16::mStr) fluorescent fusion proteins. In this cell line, *LmxFused* and *LmxULK4* were then deleted, respectively (Fig 3A-B). This showed that Δ*LmxFused* and Δ*LmxULK4* exhibit highly similar proportions of multiple distinct flagellar morphotypes and effectively phenocopy each other (Fig 3B). ~2/3 of cells in both KO cell lines not only lacked a long external flagellum (as observed in the untagged cell lines (Fig 2B, D), but the absence of RSP11::eYFP and PF16::mStr fluorescent signals at the anterior cell pole, where the flagellum normally emerges from the flagellar pocket, indicated that these cells lacked axonemes entirely (Fig 3B, v).

**Fig 3.**
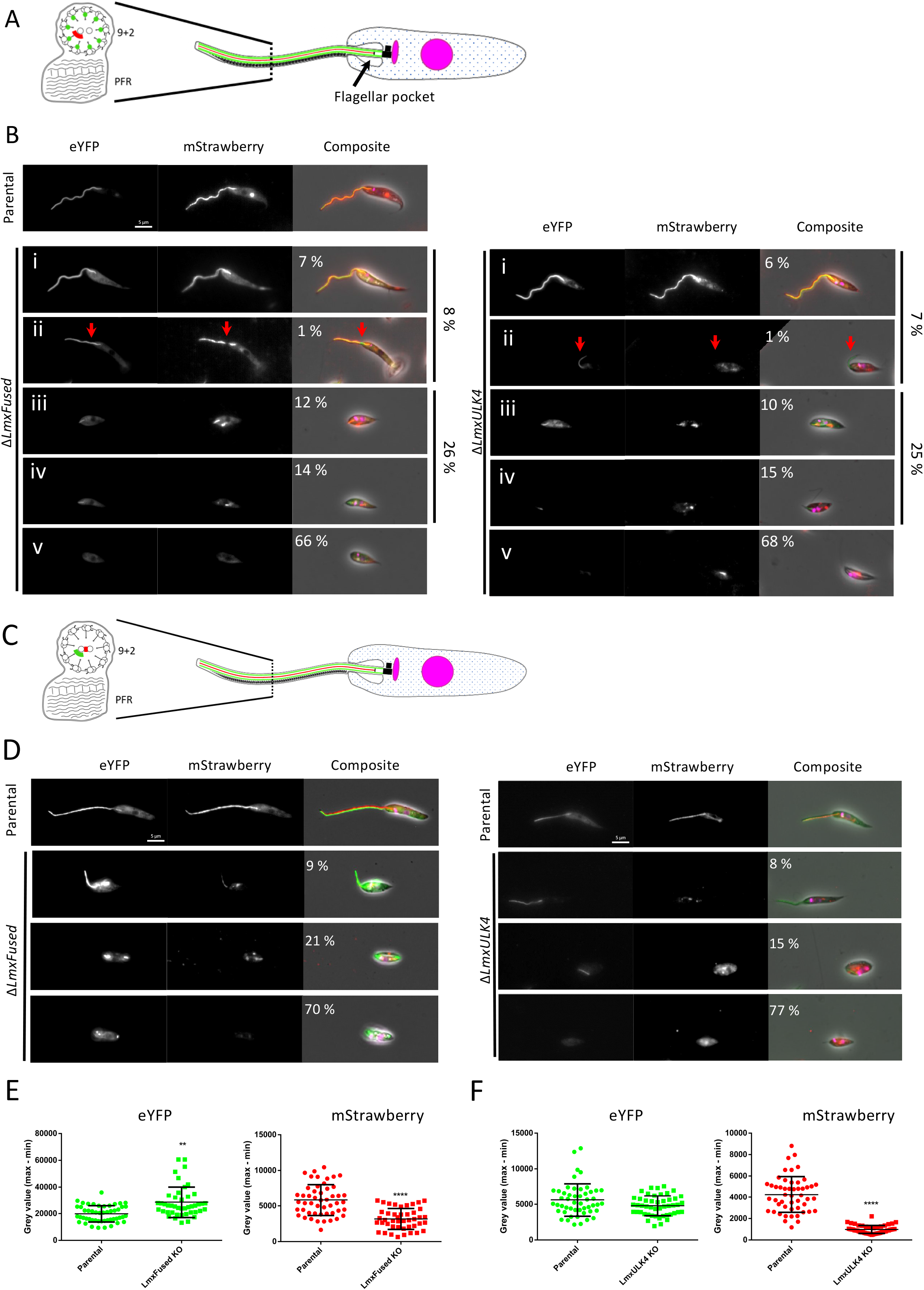
Quantification of distinct flagellar morphotypes in LmxULK4 and LmxFused KO lines. **(A)** Cartoon depicting the expected flagellar localisation of fluorescently labelled RSP11::eYFP (green) and PF16::mStr (red) within the parental cell line. **(B)** Micrographs showing an example of each distinct flagellar morphotype observed in the tagged cell line (parental) and upon deletion of *LmxFused* (left panel) or *LmxULK4* (right panel) (n ≥ 500). Flagellar morphotypes were defined as follows: (i) a long external flagellum with no apparent defect (ii) a long flagellum with reduced or non-continuous PF16::mStr signal (highlighted by red arrows), (iii) a flagellum restricted to the flagellar pocket, (iv) a flagellum restricted to flagellar pocket lacking PF16::mStr signal, or (v) no flagellar structure detected, as assessed by the absence of fluorescently labelled proteins at the anterior cell pole where the flagellum normally emerges. **(C)** Cartoon depicting the expected flagellar localisation of PF16::eYFP (green) and mStr::PF20 (red) within the parental cell line. **(D)** Micrographs showing an example of each distinct flagellar morphotype observed in the tagged cell line (parental) and upon deletion of *LmxFused* (left panel) or *LmxULK4* (right panel) (n ≥ 500). The composite images in (B and D) are overlays of the phase contrast image with the red and green fluorescence channels and the Hoechst 33342 fluorescence indicating nuclear and kinetoplast DNA. **(E-F)** Comparison of flagellar PF16::eYFP and mStr::PF20 signal intensity in the parental cell line with Δ*LmxFused* **(E)** and Δ*LmxUlk4* **(F)** cell lines. This analysis only included cells with a long external flagellum with a continuous PF16::eYFP signal.

Defects in the CP were previously reported for distinct ciliated tissues in both ULK4 and Fused/STK36 mutant mice (Liu et al., 2016; Nozawa et al., 2013; Wilson et al., 2009). Moreover, the central pair microtubule associated protein PF20 (but not PF16) has previously been shown to co-immunoprecipitate with Fused/STK36 when expressed in HEK 293T cells (Wilson et al., 2009). Examination of PF16::mStr signals (Fig 3A-B) found that a majority of cells with a long flagellum still possessed a continuous PF16 signal (Fig 3B), suggesting that the CP was still assembled in these cells, and a disrupted signal was only observed in a small percentage of flagella. To examine whether PF20 was affected by the loss of *LmxFused* and *LmxULK4*, each gene was deleted in a reporter cell line expressing PF16::eYFP / mStr::PF20 (Fig 3C-D). Focusing on the external flagella that still showed a continuous PF16::eYFP signal, we measured the intensity of fluorescence and detected a reduced signal intensity for mStr::PF20 in both Δ*LmxFused* and Δ*LmxULK4* cell lines, while no reduction of the PF16::eYFP signal was seen (Fig. 3E-F). This suggests that PF20 appears to be preferentially lost from Δ*LmxFused* and Δ*LmxULK4* axonemes. In the reciprocal experiments, KO of *LmxPF20* did not appear to impact either LmxULK4::mNG or LmxFused::mNG expression or localisation (S4 Fig).

Consistent with these findings, cross-sections through free flagella imaged by transmission electron microscopy (TEM) showed normal 9+2 microtubule arrangements and PFR structures in the majority of flagella (Fig 4A, i-iii). No obvious central pair defects were noted in any of these TEM profiles, consistent with the observed RSP11::eYFP and PF16::mStr signals in the majority of external flagella and despite the reduction of PF20 signal (Fig 3). Since fewer than 1% of cells in both Δ*LmxFused* and Δ*LmxULK4* populations showed a non-continuous or absent PF16::mStr signal (Fig 3B), the number of external flagella assessed by TEM was likely too small to capture those rare cells. We also noted that one out of the thirteen Δ*LmxFused* flagella captured by TEM displayed a 10+2 microtubule arrangement (Fig 4A, iv), indicating that a variety of ultrastructural defects may be present within sub-sets of these long external flagella.

**Fig 4.**
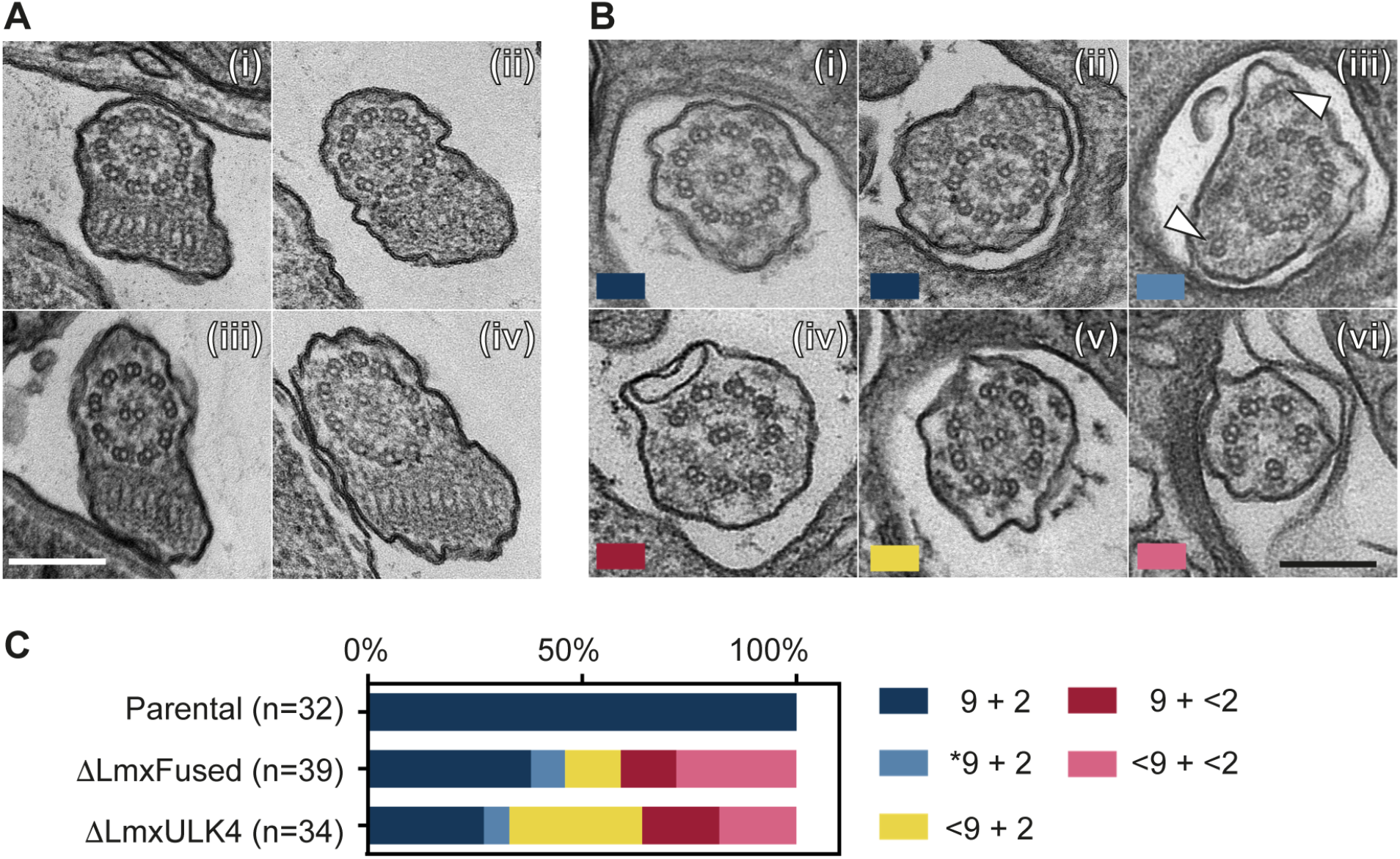
Ultrastructural defects in ΔLmxFused and ΔLmxUlk4 axonemes. Transmission electron microscopy images showing cross-sections of flagellar axonemes. **(A)** Sections through the distal part of the flagellum outside of the flagellar pocket: (i) parental, (ii) Δ*LmxULK4*, (iii, iv) Δ*LmxFused*. **(B)** Sections through the proximal part of the flagellum within the flagellar pocket show examples of normal 9 + 2 microtubule arrangements in (i) parental and (ii) Δ*LmxFused* flagella, and examples of ultrastructural defects in Δ*LmxFused* (iii-vi). Colour labels indicate categories as quantified in C. **(C)** Quantification of ultrastructural defects of axonemes within the flagellar pocket. The categories were based on the number and arrangement of microtubules (MT). 9+2 (dark blue) represents the normal configuration of 9 radially symmetric outer doublet MTs and two singlet MT; *9+2 indicates a collapse in the radial symmetry (light blue); <9+2 (yellow), 9+<2 (dark pink), and <9+<2 (light pink) indicate deviations from the expected MT numbers.

Since fluorescence microscopy data showed that ~25% of KO cells possessed a short flagellum that failed to extend significantly beyond the neck region of the flagellar pocket (Fig. 3, iii-iv), the structure of axonemes inside the flagellar pocket was examined next. Here, only 38% Δ*LmxFused* and 27% of Δ*LmxULK4* axonemes viewed by TEM showed a 9+2 MT arrangement, compared to 100% in the parental controls (Fig 4B, C). A variety of defects were noted in both central pair and outer microtubule doublet number and orientation (Fig 4B, C), including mis-positioned outer doublet MT (Fig 4B, iii), <9 doublet MTs (Fig 4B, v and vi) and absence of CP singlet MTs (Fig 4B, iv and vi). The proportion of axoneme sections lacking a CP (Fig 4C) was consistent with the absence of a PF16::mStr signal in about half of the short axonemes viewed by fluorescence microscopy (Fig 3B). To understand better where the defects originated, serial TEM sections spanning the basal-body, transition zone and axoneme regions were examined (S5 Fig A-E). Some cells only extended a few singlet or doublet microtubules from a basal body (S5 Fig B,C); such cells were likely scored as lacking axonemes entirely in Fig 3 (category v in Fig 3B). Other cells had more subtle defects, showing alteration of MT numbers or positioning (S5 Fig D,E). Taken together this suggests that all MT structures of the flagellum are sensitive to the loss of *LmxFused* and *LmxULK4* but none are strictly dependent on these proteins for their formation.

### IFT recruitment and processivity correlates with ΔLmxFused and ΔLmxULK4 flagellar morphotype

To determine whether the loss of *LmxFused* and *LmxULK4* affected intraflagellar transport, two cell lines were generated that expressed fluorescently tagged IFT proteins, mNG::IFT81/LmxM.33.0230 and mNG::IFT140/LmxM.31.0310 (components of the IFT-B and IFT-A subcomplexes, respectively (Nakayama and Katoh, 2018)). *LmxFused* and *LmxULK4* were then knocked out in these reporter lines, which presented with the typical flagellar abnormalities described above. In the minority population that still possessed a long flagellum, IFT particle numbers and velocity were measured, and the results showed that the trains migrated at a relatively normal velocity along Δ*LmxFused* and Δ*LmxULK4* flagella (S6 Fig A-N). This analysis demonstrates that in the absence of *LmxFused* and *LmxULK4* at least some cells retain the capacity for normal IFT in fully grown flagella and does not preclude the possibility that loss of *LmxFused* or *LmxULK4* KO could directly impede IFT in newly growing flagella or in the cells that fail to construct an axoneme at all. Indeed examination of the fluorescence signals showed that a minority population of Δ*LmxULK4* and Δ*LmxFused* cells didn’t have IFT signal foci in the basal body region. The mNG::IFT81 signal was absent in 6% of Δ*LmxFused* cells and in 7% of Δ*LmxULK4* cells; the mNG::IFT140 focus at the flagellum base was absent in 37% of Δ*LmxFused* cells and in 28% of Δ*LmxULK4* cells (S6 Fig O,P). Taken together these data show that the IFT behaviour in Δ*LmxFused* and Δ*LmxULK4* cells correlated with the flagellar morphotype. It remains to be tested whether this apparent reduction in IFT protein recruitment is caused by the loss of LmxFused or LmxULK4, and thus a primary cause of flagellum assembly failure, or whether reduced IFT recruitment is an indirect downstream consequence of structural defects resulting from *LmxFused* and *LmxULK4* KO loss.

### ΔLmxULK4 mutants display reduced α-tubulin acetylation, though ΔLmxFused mutants do not

A recent study has shown that RNAi mediated knockdown of *ULK4* resulted in reduced levels of acetylated α-tubulin in both primary cultured mouse neurons and brain sections (Lang et al., 2016), though that study did not examine cilia or flagella directly. Alpha-tubulin acetylation has previously been linked to increased microtubule stability and flexibility (see Janke and Montagnac, 2017; Portran et al., 2017). To test the α-tubulin acetylation levels in the *L. mexicana* parental and KO cell lines, cells were stained with antibody C3B9. This revealed a strong signal along the length of the flagellum and a cortical cell body signal, indicating detection of acetylated tubulin in the axonemal microtubules as well as the cortical cytoskeleton (Fig 5A). Knockout of *LmxULK4* resulted in a reduction in acetylated α-tubulin signal in both the *Leishmania* cell body and flagellum (Fig 5A-B). That signal was restored upon expression of an episomal add-back copy of *LmxULK4* (Fig 5A-B). However, no such reduction in acetylated α-tubulin was noted in the Δ*LmxFused* cell line (Fig 5A-B). As a control, we knocked out *Lmx*α*TAT1* (LmxM.25.1150), the ortholog of the dominant mammalian α-tubulin acetyltransferase. These cells showed a near complete loss of acetylated α-tubulin signal (Fig 5A-B). Interestingly, however these cells possessed normal long flagella (Fig 5C), and swam normally when observed in culture. Taken together, these data show that the level of α-tubulin acetylation did not correlate with the capacity to build normal motile flagella in these *Leishmania* mutants, and that the flagellar defects seen in Δ*LmxFused* and Δ*LmxULK4* cell lines were not caused by a reduction in tubulin acetylation. ULK4 may instead regulate α-tubulin acetylation in a Fused/STK36 independent manner in *Leishmania*, and perhaps mammals as well (Lang et al., 2016).

**Fig 5.**
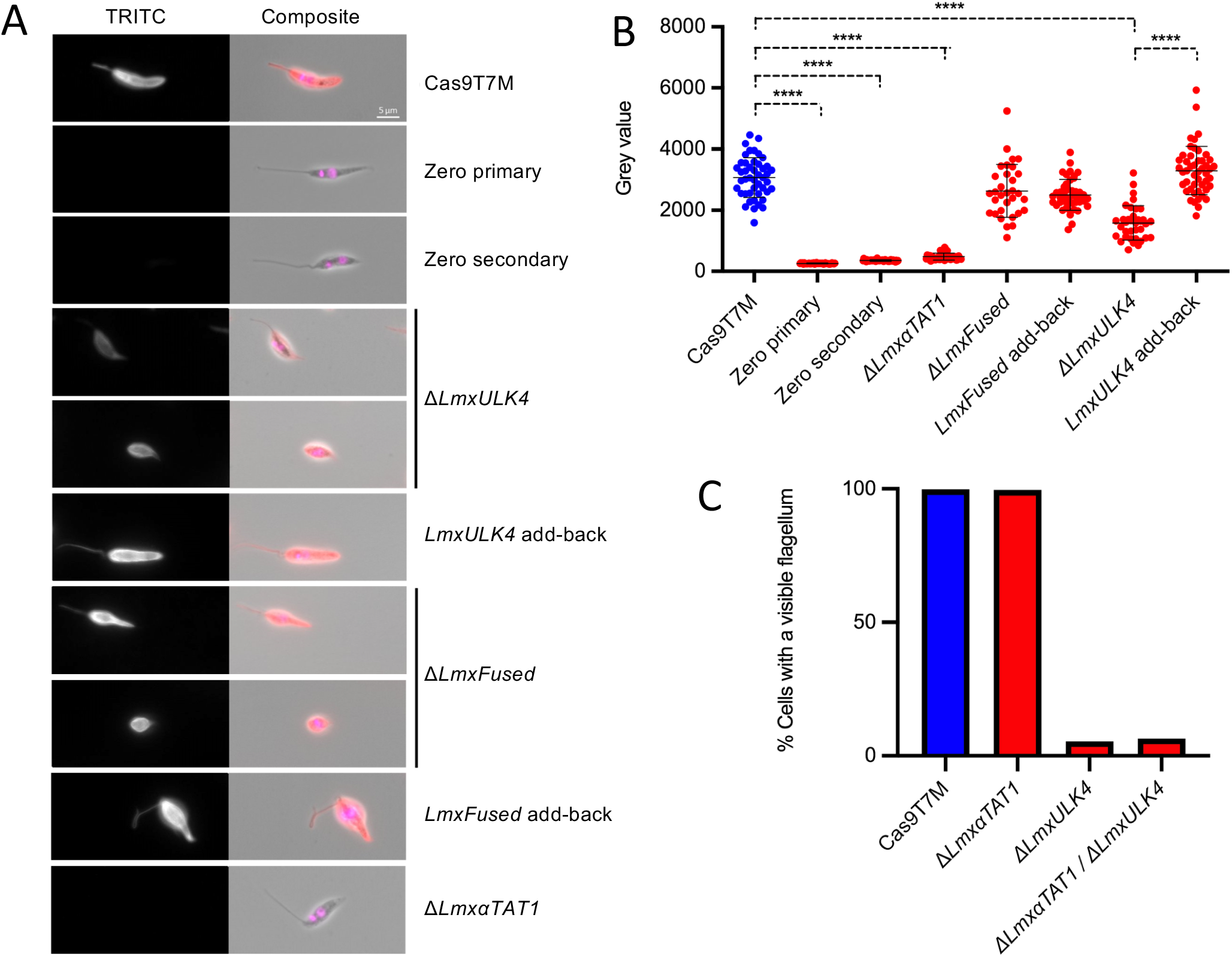
Acetylated α-tubulin levels in ΔLmxULK4 and ΔLmxFused cell lines. **(A)** Acetylated α-tubulin was detected via immunofluorescence using monoclonal antibody C3B9 and TRITC-conjugated secondary antibody. DNA was labelled with Hoechst 33342. **(B)** Comparison of flagellar TRITC signal intensity between the parental control and Δ*LmxULK4*, Δ*LmxFused* and Δ*Lmx*α*TAT1* deletion mutants (n ≥ 32 cells). For Δ*LmxULK4* and Δ*LmxFused*, cells that displayed a long external flagellum were included in the analysis. **(C)** Percentage of cells displaying a visible long external flagellum that extends beyond the flagellar pocket in the parental cell line (99.8%) populations of Δ*Lmx*α*TAT1* (99.6%) Δ*LmxULK4* (5.4%) and Δ*LmxULK4/*Δ*Lmx*α*TAT1* (6.4%) (n ≥ 500).

### Interaction between LmxULK4 and LmxFused

The similarities of the Δ*LmxFused* and Δ*LmxULK4* flagellar phenotypes individually and in combination (see Fig 2-4) suggests that *LmxFused* and *LmxULK4* may act in a common pathway. To determine whether they are in close spatial proximity, and to identify other proximal proteins or complexes, we tagged both alleles of *LmxFused* with the promiscuous *E. coli*-derived biotin ligase BirA* and preformed proximity dependent biotinylated protein identification (BioID (Roux et al., 2012)). Expression of LmxFused::BirA* had no effect on the growth of the cells in culture (S7 Fig A) and the protein showed the expected localisation (S7 Fig B). Biotinylated proteins were captured with streptavidin-coupled beads and western blot analysis confirmed the presence of the bait protein LmxFused::BirA* in the eluted fraction (S7 Fig C), which was then analysed by nano-LC/MS/MS. The parental cell line subjected to the same pull-down and MS analysis served as a control. As expected, several endogenously biotinylated proteins (putative carboxylase subunits) were identified and highly represented in both samples (supplemental data file 1). We used SINQ analysis (Trudgian et al., 2011) to test for enrichment of biotinylated proteins in the LmxFused::BirA* line and identified 125 enriched proteins (supplemental data file 1): 47 were only detected in the LmxFused::BirA* sample (supplemental data file 1 and Fig 6A) and 78 proteins showed ≥2 fold enrichment within the LmxFused::BirA* sample (supplemental data file 1 and Fig 6B). Notably, LmxULK4 was the most highly represented protein identified specifically within the LmxFused::BirA* sample after the bait protein itself (Fig 6A), suggesting that these proteins are in close proximity to each other in the cell. To test whether there was direct physical interaction between LmxFused and LmxULK4, we performed immunoprecipitation, in a cell line expressing LmxFused::eYFP and LmxULK4::MYC. Expression of fusion proteins was confirmed by fluorescence imaging (LmxFused::eYFP, S8 Fig A) and western blots of whole cell lysates (both proteins, S8 Fig B and C). Cell lysates were incubated with anti-GFP conjugated beads and protein pulldown of LmxFused::eYFP was confirmed by western blot (S8 Fig B). A control cell line expressing GFP served as a control for the pull-down and MS analyses (S Fig 8 D). The eluates were analysed by mass spectrometry (supplemental data file 2). The two highest scoring hits were a carbamoyl-phosphate synthase (LmxM.16.0590) and LmxULK4. Notably, 29 of 30 ULK4 peptides mapped specifically to the N-terminus of LmxULK4 (S8 Fig E), within the first 340 out of 1315 amino acids. Western blots probing for the C-terminal tag on LmxULK4::MYC suggested the protein was cleaved or degraded following cell lysis (S8 Fig C). The detection of peptides derived from the LmxULK4 N-terminus in IP eluates is consistent with a direct physical interaction between LmxFused and N-terminal region of LmxULK4, which also agrees with data on mammalian ULK4 interacting via its kinase domain with Fused/STK36 (Preuss et al., 2020).

**Fig 6.**
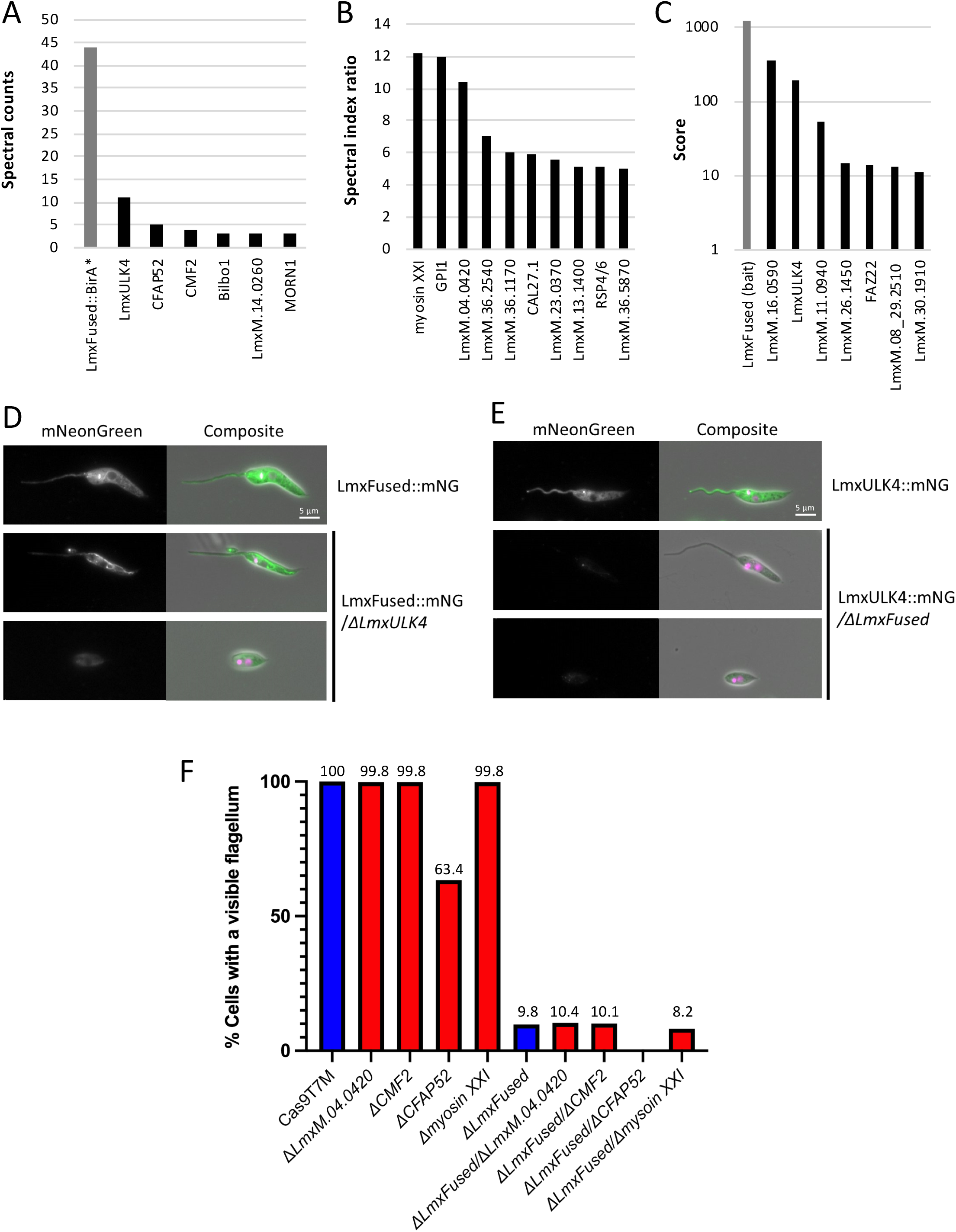
BioID and IP data support an interaction between LmxFused and LmxULK4. **(A)** Proteins that were identified by BioID only in the LmxFused::BirA* samples **(B)** Proteins identified by BioID that were ≥ 5 fold enriched within the LmxFused::BirA* samples relative to the parental control (Cas9T7M). **(C)** Proteins that co-immunoprecipitated with LmxFused::eYFP, and for which ≤1 peptides were identified in the parental or GFP control sample. **(D)** Visualisation of LmxFused::mNG in live promastigotes before and after the deletion of *LmxULK4*. **(E)** Visualisation of LmxULK4::mNG in live promastigotes before and after deletion of LmxFused. The composite images show overlays of phase contrast images with the green fluorescence channels and fluorescence of Hoechst 33342 to indicate nuclear and kinetoplast DNA. **(F)** The percentage of cells displaying a visible long external flagellum that extends beyond the flagellar pocket was quantified in promastigote populations following KO of myosin XXI, LmxM.04.0420, CFAP52 or LmxM.27.0870 (CMF2) individually, or in combination with KO of LmxFused (n ≥ 500).

Given the data presented above supporting an interaction between LmxFused and LmxULK4, we investigated the impact of gene KOs for one of these proteins on the subcellular localisation of the other protein. *LmxULK4* KO did not appear to affect LmxFused::mNG localisation (Fig 6D). In the equivalent reciprocal experiment, the signal from LmxULK4::mNG (both alleles tagged) disappeared upon deletion of *LmxFused*, suggesting that the stable expression of LmxULK4 was dependent on the presence of LmxFused (Fig 6E). These data further corroborate the KO phenocopy and together with the LmxFused BioID and Co-IP results, which both identified LmxULK4 as high-scoring interaction partner, support the hypothesis that LmxULK4 and LmxFused physically and functionally interact to contribute to the assembly of a normal motile eukaryotic flagellum.

### Interactions with other flagellar proteins

The targets of the LmxFused kinase remain unknown. BioID and Co-IP both identified additional proteins, albeit with little overlap between the two methods (supplemental data files 1 and 2). Of the 24 proteins identified in the co-immunoprecipitation with LmxFused::eYFP (≥ 3 spectral counts), 15 were also co-immunoprecipitated with the soluble GFP pulldown control sample, suggesting that these proteins directly interact with eYFP tag as opposed to LmxFused itself (supplemental data file 2). Besides LmxULK4, there were only six proteins detected with > 2 peptides in the LmxFused Co-IP sample (and < 2 peptides in the GFP pulldown and parental control samples): LmxM.16.0590 (putative carbamoyl-phosphate synthase), LmxM.11.0940 and LmxM.26.1450 (both hypothetical proteins, present in the flagellar proteome (Beneke et al., 2019)), LmxM.36.4330 (the *L. mexicana* orthologue of *T. brucei* flagellar attachment zone (FAZ) protein FAZ22 (Zhou et al., 2018)), LmxM.08_29.2510 (glycosomal i-dependent 6-phosphofructokinase) and LmxM.30.1910 (hypothetical protein) (Fig 6C). Further experiments will be required to establish which, if any, of these are genuine interaction partners of LmxFused/LmxULK4.

In addition to identifying components of a protein complex, BioID could provide information about the subcellular domains where LmxFused resides, identifying proteins that do not directly interact with LmxFused but that are in close proximity to it. Besides LmxULK4, the 124 enriched proteins included 23 that have been previously identified within the *L. mexicana* flagellar proteome (Beneke et al., 2019) and 7 that associate with the FAZ, consistent with the observed subcellular localisation pattern of LmxFused. We hypothesised that potential substrates of LmxFused, or direct interaction partners may be found amongst the most highly enriched proteins in the LmxFused::BirA* sample. We used gene KO to determine whether deletion of these genes had a similar effect as loss of Fused, and whether a dual KO modulated the Δ*LmxFused* phenotype, as assessed by quantifying the proportion of cells with a long flagellum (Fig 6F). We included in this experiment the proteins that were identified only in the LmxFused::BirA* sample with ≥4 spectral counts (LmxM.28.1110, Cilia And Flagella Associated Protein 52 (CFAP52), and LmxM.27.0870 (Component of Motile Flagellum 2 (CMF2)) and those that were highly enriched in LmxFused::BirA* sample compared to the control (spectral index ratio ≥10; LmxM.31.3870 (putative myosin XXI), LmxM.08_29.2030 (putative N-acetylglucosamyl transferase component GPI1), and LmxM.04.0420 (Tetratricopeptide repeat protein)). Loss of myosin XXI, LmxM.04.0420 and LmxM.27.0870 (CMF2) either individually or in combination with Δ*LmxFused* did not alter the phenotype of the respective parental cell line (Fig 6F). For GPI1 this could not be assessed, as we were unable to generate a null mutant. By contrast, KO of *LmxCFAP52* alone resulted in a 37% reduction in cells exhibiting long external flagella. Interestingly, the Δ*LmxFused /* Δ*LmxCFAP52* dual KO resulted in cells that lacked long external flagella entirely (Fig 6F), exacerbating the phenotype of the individual gene deletions. *Chlamydomonas* FAP52 was recently shown to function in the stabilisation of axonemal microtubules (Owa et al., 2019). One possible interpretation of the KO phenotypes observed here in *L. mexicana* is that flagellar stability is compromised to different degrees by the loss of CFAP52 and the loss of LmxFused, and when both are lost together, cells can no longer build any flagella.

### Mammalian ULK4 localises to motile cilia of mouse ependymal cells

Previous studies have provided several strong lines of evidence linking mammalian Fused/STK36 and ULK4 to motile cilia function (Liu et al., 2016; Wilson et al., 2009) and biochemical evidence indicated proximity of mammalian Fused/STK36 and ULK4 (Preuss et al., 2020). Mammalian Fused/STK36 was shown to localise along the entire length of the motile cilia (Nozawa et al., 2013) but the localization of mammalian ULK4 to motile cilia has not been assessed. We therefore transiently expressed mouse ULK4 (MmULK4) tagged with Flag-HA-mNeonGreen in the primary cell culture of mouse ependymal cells competent of forming motile cilia; aberrant axonemal structure was previously observed in this cell type of *Ulk4* hypomorph mice (Liu et al., 2016). Staining fixed cells with an antibody against acetylated tubulin as an axonemal marker, revealed that the tagged MmULK4 localised to multiple foci in the apical region of the cell, in which the cilia originated, and at distal tips of short axonemes (Fig. 7A). An accumulation, albeit with lower signal intensity, was noticeable also at the tips of a subset of longer axonemes. Finally, a weak but discernible signal was observed along each axoneme, and a diffused signal in the cytosol (Fig. 7B).

**Fig 7.**
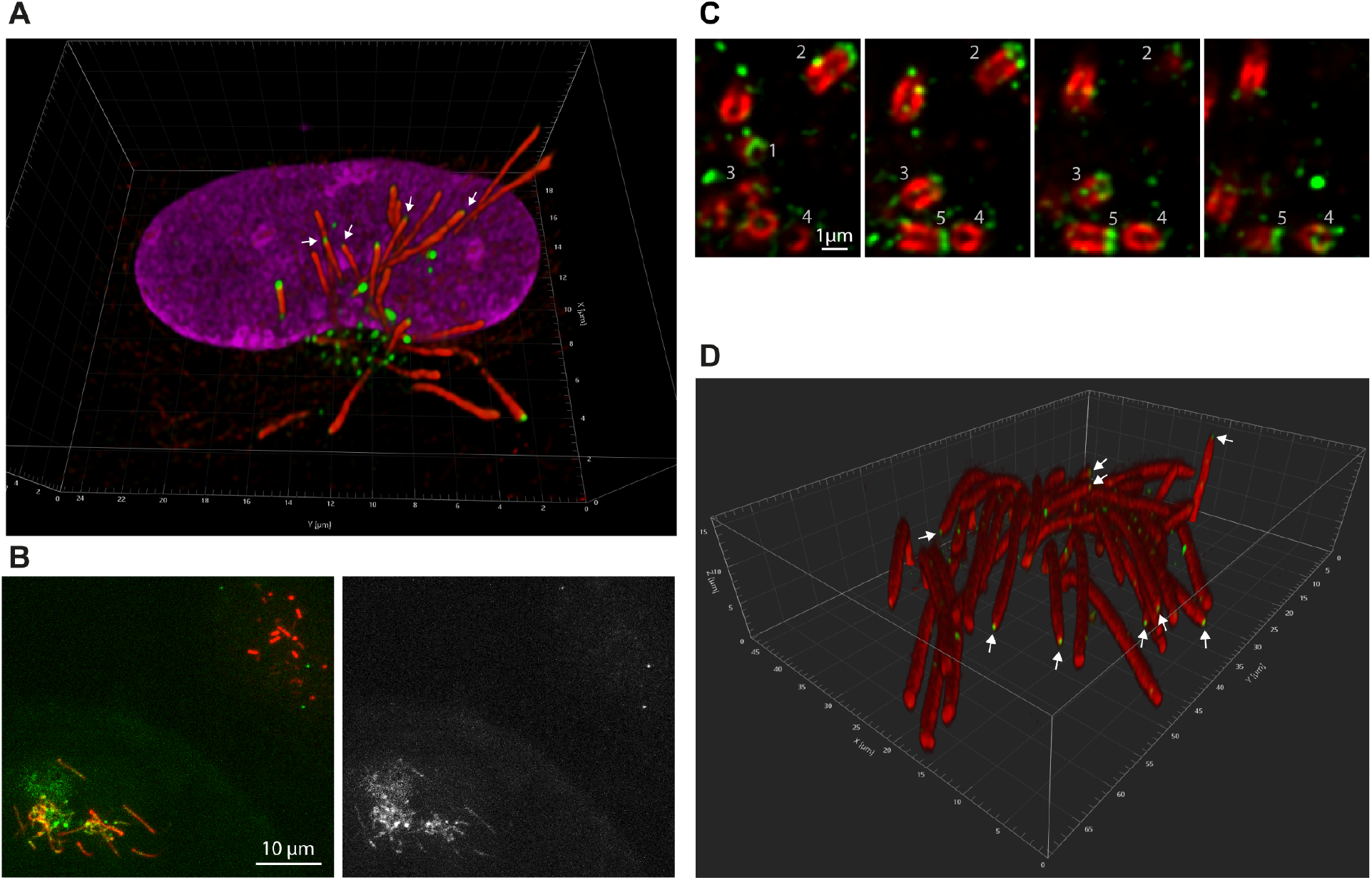
Localisation of ULK4 in mouse ependymal cilia. **(A)** Immunofluorescence staining of a multi-ciliated mouse ependymal cell. 3D reconstructed confocal z-stack using Imaris viewer (blend mode). Z-step size was 130 nm. Acetylated tubulin (labeled with C3B9) in red, Flag-HA-mNeonGreen-tagged MmUlk4 (mNG fluorescence) in green, DNA (DAPI) in magenta. Note the strong mNG signal at tips of short axonemes and the weaker signal associated with tips of certain longer axonemes (arrows). Note that under used conditions the anti-acetylated antibody does not label centrioles in immunofluorescence. **(B)** Maximum intensity projection of confocal z-stack encompassing the apical region of two multi-ciliated cells, one expressing Flag-HA-mNeonGreen-tagged MmULK4 (bottom left) and the other not (top right). To improve the signal to noise ratio, we used 4x line averaging. Raw data are presented. Left-ciliary marker Arl13b (labeled with anti-Arl13b antibody) in red, Flag-HA-mNeonGreen-tagged MmUlk4 (mNG fluorescence) in green. Right-mNG signal only. **(C)** Selected confocal planes of an apical region of an expanded multi-ciliated mouse ependymal cell, which encompasses centrioles. Individual planes are separated by 700 nm in the axial direction. Individual centrioles (labeled with C3B9; in red) are denoted with numbers. Note the Flag-HA-mNeonGreen-tagged MmUlk4 signal (anti-HA staining; in green) capping one end of each centriole. **(D)** 3D reconstruction of a confocal z-stack encompassing distal parts of cilia of an expanded multi-ciliated mouse ependymal cell. Z-step size was 100 nm and Imaris viewer (blend mode) was used for the reconstruction. Acetylated tubulin (labeled with C3B9) in red, Flag-HA-mNeonGreen-tagged MmULK4 (anti-HA signal) in green. Arrows indicate the tips of axonemes with detectable MmULK4.

We next wanted to assess whether the dot-like signals in the apical region could be associated with centrioles. As the combination of the anti-acetylated tubulin antibody and formaldehyde fixation was well suited for the visualisation of axonemes in immunofluorescence, but not for centrioles, we used expansion microscopy, which was previously shown to facilitate access of antibodies to their respective epitopes (Tillberg et al., 2016). Following the ultrastructure expansion microscopy protocol from (Gambarotto et al., 2019) we achieved about 4.7-fold expansion. Indeed, we observed that centrioles in the apical region, which did not extend axonemes, had the Flag-HA-mNeonGreen-MmULK4 associated with them, as determined by anti-HA staining (Fig. 7C). The signal was always juxtaposed distal to one end of the centriole as defined by the anti-acetylated tubulin signal (Fig. 7C). Moreover, expansion microscopy confirmed that Flag-HA-mNeonGreen-MmULK4 was present at the very tips of a subset of long axonemes (Fig. 7D).

Taken together, our data support a function for ULK4 and Fused/STK36 in motile cilia from evolutionarily distant cells, and suggest a model in which ULK4 and Fused/STK36 interact with each other to enable stable assembly of motile flagella and cilia across eukaryotic cells.

## Discussion

Here we characterised the phenotypic consequences of *Fused/STK36* and *ULK4* knockout on motile flagellum assembly in the protozoan *Leishmania mexicana*. We found that Δ*LmxULK4* and Δ*LmxFused* cells lines display equivalent flagellar assembly defects, and as such effectively phenocopy each another (Fig 2 and 3) and suggest they function within the same pathway. Intriguingly, the severity of flagellar structure defects varied between cells within knockout populations (Fig 2, 3 and 4), yet the proportion of distinct flagellar morphotypes remained consistent across multiple independent deletions of both genes and in different parental cell lines (Fig 2, 3 and S6 Fig). These data are consistent with the loss-of-function phenotypes described by Baker et al., (Baker et al., 2021) in a knockout screen of *Leishmania* kinases, and suggest the proteins are involved in regulating ciliogenesis or ciliary stability, but are not essential. In contrast, RNAi mediated knockdown of Fused and ULK4 in the related protist *T. brucei* (Jones et al., 2014; Varga et al., 2017) did not affect flagellar assembly. A small proportion of cells in which FCP6/TbFused was knocked down showed evidence of aberrant cytokinesis, which may be linked to perturbation of motility (Varga et al., 2017). The differences in the severity of phenotype between *L. mexicana* and *T. brucei* may be a result of the distinct reverse genetics methods employed (i.e. gene knockout versus knockdown). Alternatively, it could suggest that *T. brucei* possesses an unknown compensatory failsafe mechanism that results in increased flagellar assembly robustness when faced with Fused or ULK4 functional perturbation. A third possibility is that there is some divergence between Fused and ULK4 functions in *T. brucei* and *L. mexicana* that could be linked to the differences in cellular architecture. *T. brucei* flagella have extensive structural connections to the cell body and the newly assembling flagellum is linked to the existing flagellum through the FC structure where Fused or ULK4 were localised. The *L. mexicana* flagellum is only connected in its proximal part to the to the cell body, via a FAZ structure (Wheeler et al., 2016) and lacks a recognisable FC structure.

In addition to the phenocopy of Δ*LmxFused* and Δ*LmxULK4* described above, we show that LmxFused and LmxULK4 are in close proximity to one another (LmxFused::BirA* mediated BioID; Supplemental data file 1; Fig 6A), they co-immunoprecipitate (via LmxFused::eYFP pull down; supplemental data file 2; Fig 6C), and the stable expression of LmxULK4::mNG is dependent upon the presence of *LmxFused* (Fig. 6E). Together these data suggest that LmxULK4 and LmxFused function as part of the same pathway to mediate normal motile flagellum assembly or maintenance in *Leishmania*. Although Fused/*STK36* and ULK4 have not yet been shown to directly interact in mammals, both *Fused/STK36* and *ULK4* mutants are associated with a range of motile cilia assembly defects in distinct mouse tissues (Liu et al., 2016; Nozawa et al., 2013; Vogel et al., 2012; Wilson et al., 2009). Mammalian Fused/STK36 was previously shown to localise to the motile cilium (Nozawa et al., 2013) and we now demonstrated localization of MmULK4 to motile cilia of ependymal cells (Figure 7). Its pattern is indeed reminiscent of the pattern of LmxULK4; MmULK4 localises along the cilium with enrichment at the distal tip of the axoneme. Moreover, the accumulation at tips of nascent cilia resembles the situation in *T. brucei*, where the ortholog FCP5/TbULK4 localises to the flagella connector present at the tip of growing flagella (Varga et al., 2017). These finding point to a remarkable conservation of localization of the protein in evolution and make *Leishmania* a tractable model to dissect the mechanisms by which Fused/STK36 and ULK4 contribute to stable flagellum assembly. Interestingly, mammalian ULK4 (tagged with BirA*) has recently been shown to be in close proximity to Fused/STK36 in modified HEK 293T cells (Preuss et al., 2020), supporting the conservation of an evolutionarily ancient Fused/STK36 and ULK4 interaction that likely contributed to the assembly of motile flagella/cilia in the last common ancestor of *Leishmania* and mammals.

To date, relatively little is known about the mechanism through which either protein contribute to flagellar assembly. However, the mouse Fused/STK36 orthologue has been shown to co-immunoprecipitate with the flagella/cilia associated proteins PF20, KIF27 and PCDP1 when expressed in a HEK 293T cell line (Nozawa et al., 2013; Wilson et al., 2009). Of these three established Fused interactors, only the central pair associated protein PF20 has an obvious ortholog in *L. mexicana*. Although LmxPF20 was not shown here to be in close proximity to LmxFused::BirA*, or co-immunoprecipitate with LmxFused::eYFP, we did find that Δ*LmxULK4* and Δ*LmxFused* cells exhibited reduced levels of LmxPF20 in their long external flagella, where present (Fig 3). When considered alongside the other central pair defects noted in Δ*LmxULK4* and Δ*LmxFused* (Fig 3B and 4), these data support the idea that both ULK4 and Fused/STK36 contribute to normal central pair microtubule assembly or maintenance as proposed for mammalian cilia (Liu et al., 2016; Nozawa et al., 2013; Wilson et al., 2009). In some of those mammalian cilia the loss of ULK4 and Fused affected the integrity of the axoneme more widely, however, and in *Leishmania*, the disruption of the outer doublet microtubule arrangements and reduction of flagellar length were the dominant features of the mutant cells.

In addition to LmxULK4, several flagellar proteins were identified within our LmxFused::BirA* BioID dataset (see supplemental data file 1) as candidate LmxFused interactors. It is interesting to note that CFAP52 and CMF2 were the third and fourth most highly represented proteins identified specifically within the LmxFused::BirA* sample, after only LmxULK4 and the bait proteins itself (Fig 6). Homologs of both CFAP52 (FAP52) and CMF2 (RIB72) in *Chlamydomonas* are microtubule inner proteins (MIPs) that are bound to inner lumen wall of B and A tubules of the axonemal outer microtubule doublets, respectively (Ma et al., 2019). Both proteins contribute to enhanced microtubule stability against mechanical stress/damage (Owa et al., 2019; Stoddard et al., 2018), and human patients lacking CFAP52 (also known as WDR16) display laterality abnormalities associated with motile cilia dysfunction (e.g. *situs inversus totalis* (Ta-Shma et al., 2015)). The identification of these proteins in our BioID dataset suggests that LmxFused::BirA* may directly or indirectly interact with one or multiple MIPs to promote microtubule stabilisation. In support of this hypothesis, we have established a genetic interaction between *LmxCFAP52* and *LmxFused*, where Δ*LmxFused* / Δ*LmxCFAP52* cells display an exacerbated flagellar assembly defect relative to Δ*LmxFused* alone (they fail to build any long external flagella; see Fig 6F). FAP52 has been described as an interaction hub for a larger MIP network that includes inner junction proteins and also crosses the connection between protofilaments A12 and A13 to interact with the N-terminus of RIB72 in the A tubule (Khalifa et al., 2020; Ma et al., 2019). Individual components of this MIP network can act synergistically to promote microtubule stability (e.g. FAP52, FAP45 and FAP20 (Owa et al., 2019)), perhaps warranting the examination of further potential genetic interactions between LmxFused and other MIPs. Though not yet proven, LmxFused could interact with MIPs either prior to their incorporation into the microtubule, via diffusion along the lumen accessed at the microtubule extremities (though the rate of intra-luminal diffusion may be limiting; (Janke and Montagnac, 2017)), or through gaining luminal access at random breakage points to promote microtubule repair. Indeed, it is possible that the diversity of flagellar morphotypes described here are in part a result of the stochastic nature of microtubule destabilisation events that may occur at a higher frequency and/or remain unrepaired in the absence of LmxFused and LmxULK4 associated signalling pathways. It is worth noting that CFAP52 has also been localised to the mouse sperm manchette (Tapia Contreras and Hoyer-Fender, 2020), another microtubular structure that was adversely affected upon the loss of Fused (Nozawa et al., 2014).

It should be noted that, apart from LmxULK4, none of the flagellar proteins identified via BioID also co-immunoprecipitated with LmxFused::eYFP. This discrepancy is likely due to the fact that our IP protocol was biased towards the identification of soluble proteins, as we primarily attempted to confirm the interaction between LmxFused and LmxULK4, both of which appear to be detergent extractable (see S1 Fig). Future efforts to confirm any candidate LmxFused protein kinase substrates identified here could be aided by the generation of a suitable engineered ATP-analogue specific LmxFused kinase (Romano et al., 2017).

Though not directly tested here, ULK4 is predicted to be a pseudokinase (Fig 1B), similar to the metazoan ULK4 proteins. Interestingly, ULK4 in plants and other protists still possess the canonical catalytically relevant residues and it appears that catalytic activity may have been lost multiple times in evolution (Preuss et al., 2020). LmxFused is predicted to be an active kinase. In *Drosophila* hedgehog signalling Fused displays both catalytic and non-catalytic functions (see Maloverja & Piirsoo, (2012) for a review). By contrast, attempts to demonstrate mammalian Fused kinase activity *in vitro* were unsuccessful (Murone et al., 2000; Wilson et al., 2009)), leading to speculation that the mammalian Fused orthologue may not function as an active kinase (Maloverja & Piirsoo, 2012). However, given the conservation of canonical catalytically relevant amino acid residues within the active site of these proteins (Fig 1B), and that our results suggest that Fused and ULK4 may function together as a part of the same pathway, it may be interesting to test for mammalian Fused kinase activity in the presence of ULK4.

How might ULK4 affect the activity of Fused? Deletions of LmxULK4 KO or LmxFused KO result in the same phenotype with respect to flagellar morphotype (Fig. 2, 3 and 4) In *Leishmania* the dual KO cell line (Δ*LmxULK4* / Δ*LmxFused*) appear phenotypically indistinguishable from either Δ*LmxULK4* or Δ*LmxFused* (Fig. 2-4). This argues against an inhibitory function of ULK4. Instead ULK4 may stimulate Fused kinase activity in some way; either through the allosteric activation of Fused kinase activity, or functioning as a scaffold to facilitate Fused-substrate interactions which may not otherwise occur at a sufficient rate via diffusion alone (see Figure 8). In *Drosophila* (which lacks an ULK4 orthologue) Fused homodimerization proceeds trans auto-phosphorylation of the activation loop to promote kinase activity and normal hedgehog signalling (Shi et al., 2011; Zhang et al., 2011). It therefore seems likely that *Leishmania* orthologue may similarly require both dimerization and activation loop phosphorylation (common protein kinase activation steps) to facilitate LmxFused kinase activity. Indeed, the Fused orthologue in mice appears to oligomerise (Fused proteins tagged with alternate epitopes exhibit co-immunoprecipitation (Nozawa et al., 2013)) supporting the potential for Fused dimerisation (if not multimerization) in mammals as well. Conversely, given the precedent for established pseudokinase-kinase interactions and heterodimers, where the pseudokinase interacting partner regulates the activity of the bona fide catalytic kinase (see (Shaw et al., 2014)) for a review), it is also possible that LmxFused forms a heterodimer with LmxULK4. Shaw et al. (Shaw et al., 2014) hypothesise that pseudokinase-kinase heterodimerization may represent a common phenomenon that facilitates cis autophosphorylation and precedes kinase activation.

**Fig 8.**
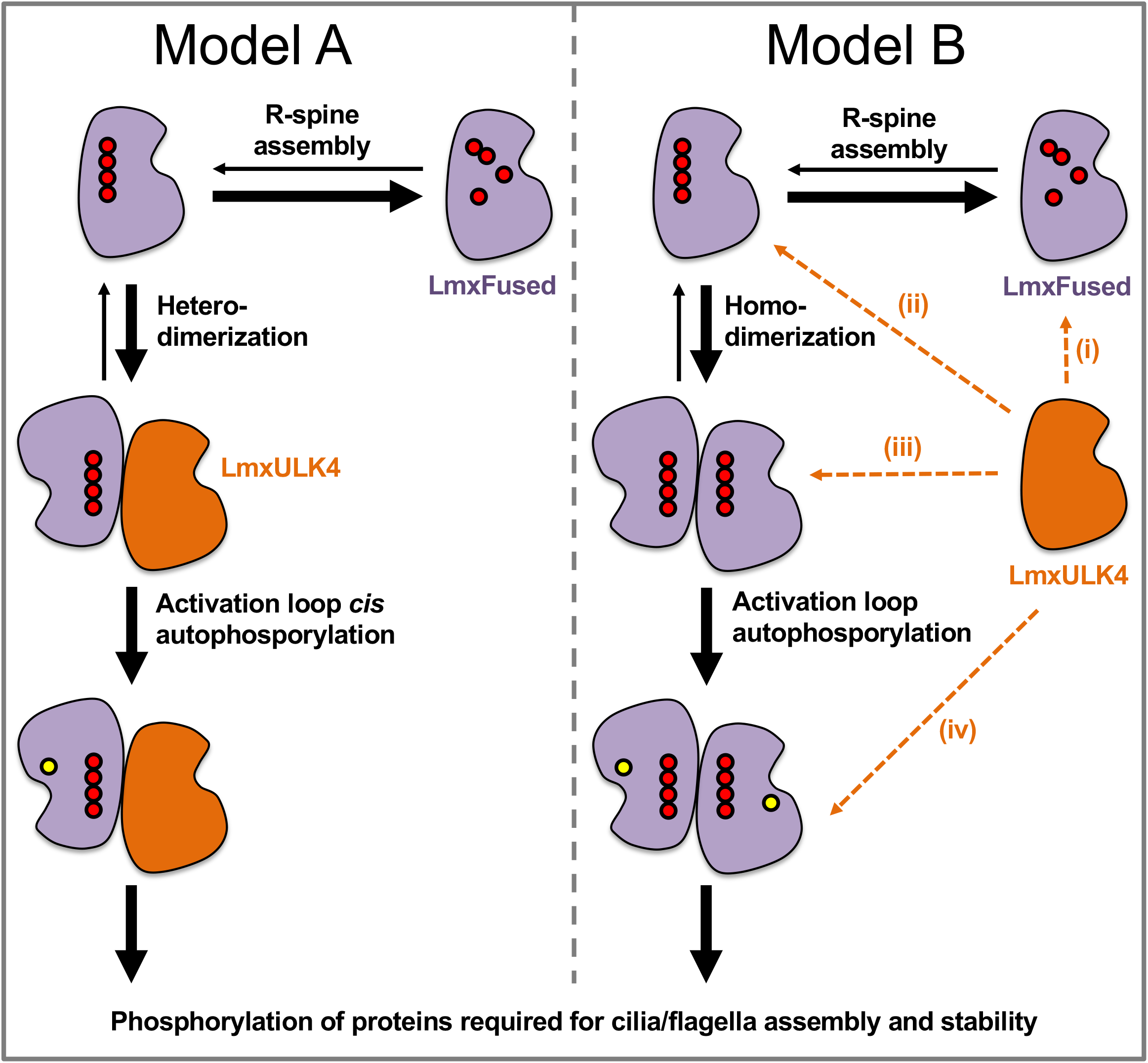
Models for possible interactions between ULK4 and Fused/STK36. In Model A, LmxFused forms a heterodimer with LmxULK4. Heterodimerization promotes *cis* autophosphorylation of the LmxFused activation loop to contribute to flagellum assembly or microtubule stabilisation. In Model B, LmxFused forms a homodimer. In this context it is not yet clear if LmxULK4 interacts with LmxFused to promote; (i) assembly of the regulatory spine (R-spine), which is a key feature of activated kinases, (ii) homodimerization, (iii) activation loop phosphorylation and/or (iv) the interaction of LmxFused with downstream targets. Stacked red circles represent the formation of the R-spine. Yellow circles represent activation loop phosphorylation.

While this study focused on the role of Fused/STK36 and ULK4 with respect to motile flagellum construction and maintenance, this interaction likely has functional relevance beyond motile cilia and flagella structure and function. In support of this idea, and as highlighted above, ULK4 was shown to be in close proximity to both Fused and several other microtubule associated proteins in HEK 293T cells (Preuss et al., 2020); a cell line which lacks motile cilia. Both proteins also localise to the same layer of the *T. brucei* flagella connector (Varga et al., 2017), though their function within this insoluble mobile transmembrane structure is not yet known. Additionally, although not yet proven experimentally, it has recently been hypothesised that the orthologous proteins of Fused (TIO) and ULK4 (RUK) may cooperate to regulate microtubule depolymerisation in the lagging edge of the cytokinesis/cell plate associated plant phragmoplast (Smertenko et al., 2018). The non-ciliated species *Dictyostelium* has a Fused ortholog called *tsunami*, which localises to microtubules and *tsunami* mutants have a chemotaxis and polarisation defect (Tang et al., 2008). This suggest that interaction with microtubules is an ancestral function of Fused kinases and the interaction between Fused/STK36 and ULK4 appears to represent an evolutionary ancient partnership that likely contributes to the regulation of microtubule stability or microtubule associated signalling pathways in diverse cellular contexts in many eukaryotic organisms.

## Materials and Methods

### Cell culture and genetic modifications

*L. mexicana* promastigotes (WHO strain MNYC/BZ/62/M379) were cultured at 28°C in M199 medium (Life Technologies) supplemented with 2.2 g/L NaHCO_3_, 0.005% haemin, 40 mM HEPES and 10% FCS, pH 7.4 (complete M199). Gene tagging and KO cell lines were generated using CRISPR-Cas9 in the *L. mex* Cas9 T7 cell line (stabilate Cas9 T7 M), as described by Beneke et al. (Beneke et al., 2017) using primer sequences retrieved from www.LeishGEdit.net (see Beneke & Gluenz (2019)), unless specified otherwise. Gene KO was confirmed via diagnostic PCR (see primers included in supplemental data file 3) used to assess presence or absence of the targeted open reading frame (Beneke & Gluenz, 2019). Mouse ependymal cell culture was established as described previously (Delgehyr et al. 2015). Briefly, the brain of a P1 mouse (C57BI/6J) was dissected. Isolated lateral ventricles were mechanically and enzymatically homogenised, and the culture was expanded. Cells were seeded into a poly-L-lysine (Sigma, P1524) coated 8-well microscopy chamber (Ibidi, 80807) for immunofluorescence assay or onto a poly-L-lysine coated Ø12 mm microscopy coverslip (Marienfeld, 117520) placed in 24-well plate for expansion microscopy. A plasmid coding for expression of Flag-HA-mNeonGreen-tagged MmULK4 (cDNA: Origene, MR217918; mNeonGreen (mNG) was provided by Allele Biotechnology and Pharmaceuticals (Shaner et al. 2013)) under constitutively active CMV promoter was transfected to the seeded cells using Lipofectamine 3000 (Invitrogen, L3000001) following manufacturer instructions. On the following day, the cells were washed twice with DMEM (Sigma-Aldrich, D6429) supplemented with 100 U/ml of penicillin (BB Pharma) and 100 μg/ml of streptomycin (Sigma-Aldrich, S9137). The cells were incubated in the serum-free medium for 8 days to induce differentiation and ciliogenesis.

### Whole genome sequencing

For the original Δ*LmxFused* and Δ*LmxULK4* cell lines, complete gene KO was also confirmed via whole genome sequencing, as described in (Beneke et al., 2019). Briefly, DNA from mutants was prepared using Illumina TruSeq Nano DNA library kit and resulting libraries were quantified using NEBNext library Quant kit. Library size was determined using Agilent High Sensitivity DNA kit on a 2100 Bioanalyzer instrument. The quantified library was multiplexed with project unrelated libraries, spiked with 1% PhiX DNA and sequenced on a NextSeq 550 (1.8 pM loading concentration). Sequencing was performed in paired-end sequencing mode (2×75 cycles, 6 and 8 cycles index read) using a NextSeq 500/550 High Output Kit v2.5 (150 cycles) following the manufactures instructions. Genome samples were de-multiplexed using bcl2fastq (Illumina), assembled using Burrow-Wheeler Aligner (Li and Durbin, 2009), sorted and indexed using Samtools (Li et al., 2009), and viewed on IGV viewer (Robinson et al., 2011).

### Episomal addback

The open reading frame of *LmxFused* was amplified from genomic DNA using primers

F: 5′-TTAGCAACTAGTATGCTTGTGACCATGGAGGACTACC-3′ and

R: 5′-TTAGCAGAATTCTCAGAGGTCCTCCTCGCTAATGAGCTTTTGCTCCAGGTCTTCTTCGCTGATCAGCTTCTGTTCACCAAGTCGATCGACGAGGTTGC-3′.

The resultant PCR products were cloned into pTadd (Beneke et al., 2017), using *Spe* I and *Eco* RI restriction sites. The *LmxULK4* addback plasmid was similarly generated using primers

F: 5′-TTAGCAACTAGTATGAACAACTATGTGCTTAATGACGAGATC-3′ and

R: 5′-TTTTCAATTGTCAGAGGTCCTCCTCGCTAATGAGCTTTTGCTCCAGGTCTTCTTCGC TGATCAGCTTCTGTTCGGAGAGTTTCTTCAGAATCTTGTTCGC-3′, and *Spe* I and *Mfe* I restriction sites. 1 µg of circular plasmid was transfected as described previously (Beneke et al., 2017) to allow episomal expression of LmxFused::2xMYC and LmxULK4::2xMYC. Drug-resistant cells were selected using 25 µg/ml phleomycin.

### Live cell microscopy of Leishmania

Cells were harvested from log phase culture by centrifugation (800*g*, 5 minutes), washed once in 1 ml PBS (containing, if required, 10 μg/ml Hoechst 33342), resuspended in 10 μl PBS and imaged live, whilst adhered to a poly-lysine coated glass slide. Cells were imaged on either a Zeiss Axioimager.Z2 microscope with a 63× numerical aperture (NA) 1.40 oil immersion objective and a Hamamatsu ORCA-Flash4.0 camera, or a 63× NA 1.4 objective lens on a DM5500 B microscope (Leica Microsystems) with a Neo sCMOS camera (Andor Technology) at ambient temperature (~25–28°C). Micrographs were processed using Fiji (Schindelin et al., 2012).

### Quantification of IFT train migration and parasite motility analysis

IFT fluorescence video micrographs were captured using methods based on those employed by Wheeler at al. (2015), with 600 frames captured per video at 50 ms fluorescence exposure per frame (~95 ms delay between frames). IFT particle velocity, intensity and number were quantified using the publicly available software *KymographClear* (ImageJ macro toolset) and *KymographDirect* (see (Mangeol et al., 2016)), including correction for both background signal and signal bleaching. Short particle tracks were removed from the analyses where necessary to avoid pseudo-replication (multiple measurements of an individual particle). This analysis was restricted to cells that possessed a flagellum that extended significantly beyond the flagellar pocket. Promastigote motility assays using darkfield video microscopy were performed using methods described by (Wheeler, 2017), including all three modifications outlined by Beneke et al. (Beneke et al., 2019).

### Preparation of cells for immunofluorescence

*L. mexicana* were harvested from log phase culture by centrifugation (800g, 5 minutes), washed twice in 1 ml PBS and resuspended in PBS at ~3×10^7^ cells/ml. Cells were allowed to settle onto a glass slide for 30 minutes in a humid chamber and fixed in cold methanol (20 minutes at −20°C). Samples were rehydrated and washed twice in PBS before antibody labelling. Immunofluorescence assays of mouse cells was performed directly in the 8-well microscopy chamber at room temperature. Cells were washed twice with PBS and fixed with 4% formaldehyde (Sigma, F8775) in PBS for 10 minutes. The cells were subsequently washed with PBS and permeabilised with 0.5% Triton X-100 (Roth, R30512) in PBS for 5 minutes, followed by another PBS wash. The permeabilised cells were blocked for 15 minutes in a blocking buffer (2% bovine serum albumin (Roth, 8076.2), 0.1% Triton X-100 in PBS), followed by PBS wash.

### Antibody labelling and immunofluorescence imaging

The primary antibody C3B9 (mouse monoclonal IgG2b, (Woods et al., 1989) was used for staining acetylated tubulin. The fixed *Leishmania* were incubated with C3B9 diluted 1:10 in PBS for 1 hour in a humid chamber. The slides were then washed three times (5 minutes per wash) in PBS. Samples were incubated in TRITC-conjugated goat anti-mouse secondary antibody (115-025-146, Jackson ImmunoResearch Laboratories); 1:200 dilution in PBS plus 5% goat serum) for 1 hour, and again washed three times (5 minutes per wash) in PBS. Slides were mounted in 90% glycerol supplemented with 25 mg/ml 1,4-diazabicyclo[2.2.2]octane (DABCO) and 500 ng/µl diamidino-2-phenylindole (DAPI), pH 8.6, and imaged via fluorescence light microscopy as described above for live cell microscopy of *Leishmania*.

The fixed mouse cells were incubated with C3B9 diluted 1:50 in blocking buffer for 60 minutes. Next, the cells were thoroughly washed with PBS and stained for 30 minutes with Alexa Fluor 647-conjugated goat anti-mouse secondary antibody (A21235, Invitrogen) diluted 1:1000 in blocking buffer. Alternatively, the cells were stained with the primary antibody recognizing the ciliary protein Arl13b (17711-1, Proteintech), which was diluted 1:1000 in blocking buffer, followed by staining with Cy5-conjugated goat anti-rabbit (A10523, Invitrogen) secondary antibody diluted 1:1000 in blocking buffer. The labelled cells were subsequently washed with PBS and stained with 1 μg/ml DAPI in PBS for 5 minutes. Finally, the cells were washed with PBS, covered with 120 μl of 90% glycerol supplemented with DABCO, and stored at 4°C until imaging.

Mouse samples were imaged using a Leica TCS SP8 confocal microscope using an HC PL apochromatic ×63/1.40 oil objective. A fluorescence signal was detected using a combination of photomultiplier and hybrid detectors. Z-stacks were deconvolved with Huygens Professional v. 21.04 using deconvolution express mode (Scientific Volume Imaging, The Netherlands, http://svi.nl). Final 3D visualisation was rendered in Imaris viewer 9.7.2 (Bitplane). Selected confocal planes were processed using Fiji (Schindelin et al., 2012).

### Expansion microscopy

The specimen preparation was based on the U-ExM protocol (Gambarotto et al., 2019) with a few modifications. Briefly, cells were fixed overnight in 4% formaldehyde and 4% acrylamide (Sigma, A8887) in PBS at room temperature. For gelation, the cells were incubated for 30 minutes at 37°C in a humidified incubator in 50 µl of monomer solution containing 19% sodium acrylate (Sigma, 408220), 10% acrylamide, 0.1% N, N’-methylenebisacrylamide (Sigma, M7256), 0.5% N, N, N’, N’-tetramethylethylenediamine (Sigma, T9281) and 0.5% ammonium persulfate (Thermo Scientific, 17874). After denaturation (60 minutes at 95°C in denaturation buffer composed of 50 mM Tris, 200 mM NaCl, and 200 mM sodium dodecyl sulfate in ddH_2_O; pH 9.0) and expansion of the gel by incubation in ultrapure water, 1×1 cm piece of the gel was cut and incubated overnight with primary antibodies (C3B9 (Woods et al., 1989) diluted 1:10, and an anti-HA tag antibody (Cell Signaling Technology, 3724S) diluted 1:500 in 2% bovine serum albumin in PBS). After 3 × 20 minute washes with ultrapure water, the gel was stained overnight with secondary antibodies (Alexa Fluor 488 anti-mouse (Invitrogen, A11001) diluted 1:500, and Alexa Fluor 555 anti-rabbit (Invitrogen, A21428) diluted 1:500 in 2% bovine serum albumin in PBS). Finally, the piece of gel was washed 3 × 20 minutes and stored in ultrapure water until imaging. The expanded gel was placed on a poly-L-lysine coated glass-bottom dish (Cellvis, D35-20-1.5-N) and imaged as described above.

### Transmission electron microscopy

Cells were prepared with a chemical fixation protocol similar to that outlined by Hoog et al. (Hoog et al., 2010). Briefly, cells were fixed with 2.5% glutaraldehyde and 4% paraformaldehyde in complete M199 culture medium for 2 hours at room temperature. Fixed cells were washed six times for 10 minutes in 0.1 M piperazine-N,N′-bis(2-ethanesulfonic acid) buffer (PIPES, pH 7.2), with the forth wash supplemented with 50 mM glycine. Cells were embedded in 3% low-melting-point agar and incubated in 1% osmium tetroxide and 1.5% potassium ferrocyanide in 0.1 M PIPES buffer rotating in darkness at 4°C for 1 hour. Samples were then washed five times with ddH_2_O (5 minutes per wash), and stained with 0.5% uranyl acetate at 4°C overnight in darkness. Samples were dehydrated, embedded in epoxy resin, sectioned and on section stained as described previously (Hoog et al., 2010). Electron micrographs were captured on a Tecnai 12 TEM (FEI) with an Ultrascan 1000 CCD camera (Gatan), and processed using Fiji (Schindelin et al., 2012).

### Biotin ligase mediated proximity labelling

A cell line expressing LmxFused::BirA* was generated by tagging both *LmxFused* alleles with pPLOT-BirA*::3xmyc (Beneke et al., 2017), and selected on 20 µg/ml puromycin. Clonal cell lines were selected by limiting dilution and the modification of both alleles was confirmed via endpoint PCR (using primers that spanned the inserted BirA* tag encoding region; F: 5’-CGGGGCACTGAGCAACTTTGT-3’, R: 5’-CACCAGTGGGCAGTGTGAGC-3’). Biotinylated proteins were captured using the XL-BioID protocol, which includes a cross-linking step prior to affinity capture (Geoghegan et al., 2021): Log phase promastigote cultures (seeded at 2×10^6^ cell/ml) were allowed to grow in complete M199 supplemented with 150 μM biotin (B-4639, Sigma-Aldrich) for 18-24 hours. 4×10^8^ parasites were harvested via centrifugation (800*g* for 5 minutes), washed twice with PBS, and cross-linked with 1 mM dithiobis(succinimidyl propionate) (DSP) in 10 ml PBS plus 5% dimethyl sulfoxide (DMSO) for 10 minutes at 28°C. 1M Tris pH 7.5 was added for 5 minutes at room temperature to quench, and parasites were pelleted via centrifugation (1200*g*, 3 minutes). All subsequent steps were performed on ice or at 4°C. The supernatant was discarded and parasites lysed with sterile RIPA buffer plus protease and phosphatase inhibitors (1% NP-40, 0.5% sodium deoxycholate, 0.1% SDS, 50 mM Tris pH 7.5, 125 mM NaCl, 0.1 mM EDTA, 0.1 mM Phenylmethanesulfonyl fluoride (PMSF), 1µg/ml Pepstatin A, 1 µM trans-Epoxysuccinyl-L-leucylamido(4-guanidino)butane (E-64), 0.4 mM 1-10 phenanthroline, 1 tablet/ml cOmplete™ inhibitor cocktail (Roche), 1x PhosSTOP™ (Roche) and 1x Halt™ Proteases inhibitors single-use cocktail (Thermo Scientific)), and sonicated (three times for 5 seconds at amplitude 16 on a MSE Soniprep 150). Benzonase (BaseMuncher; 0.5 units/μl) was added to digest chromatin for 1 hour and the lysate was centrifuged at 10’000*g* for 10 minutes at 4°C. The supernatant was added to 1 mg MagResyn® streptavidin beads (MR-STV002, Resyn Biosciences); note that these beads were washed twice with 1 ml of RIPA buffer before use and rotated overnight to enable biotinylated protein binding. The beads were then washed sequentially in 500 μl of RIPA buffer (4 washes in total), followed by individual washes in 4M urea, 6M urea and 1M KCl. The beads were resuspended in 50mM TEAB (triethylammonium bicarbonate). Samples capturing an equivalent fraction of the input material were collected at multiple points during the protocol, and SDS-PAGE and immunoblots were performed by standard methods using a mouse monoclonal anti-MYC antibody (clone 4A6, 05-724, Merck).

### Mass spectrometry for BioID

Sample preparation and analysis was performed by the Advanced Proteomics Facility (Department of Biochemistry, University of Oxford) as follows: On-bead protein sample digests were performed according to the filter-aided sample preparation procedure described in (Wisniewski et al., 2009). Peptides were analysed by nano-liquid chromatography tandem mass spectrometry (nano-LC/MS/MS) on an Orbitrap Q Exactive mass spectrometer (Thermo Scientific) using Higher-energy Collisional Dissociation (HCD) fragmentation. In brief, peptides were loaded on a C18 PepMap100 pre-column (300 µm i.d. x 5 mm, 100Å (Thermo Fisher Scientific)) at a flow rate of 12 μl/min in 100% buffer A (0.1% formic acid in H_2_0). Peptides were then transferred to an in-house packed analytical column heated at 45°C (50 cm, 75 µm i.d. packed with ReproSil-Pur 120 C18-AQ, 1.9 µm, 120 Å) and separated using a 60 minute gradient from 15 to 35% buffer B (0.1% formic acid in acetonitrile) at a flow rate of 200 nl/min. Q Exactive survey scans were acquired at 70,000 resolution to a scan range from 350 to 1500 m/z, automatic gain control target 3e6, maximum injection time 50 ms. The mass spectrometer was operated in a data-dependent mode to automatically switch between MS and MS/MS. The 10 most intense precursor ions were submitted to HCD fragmentation using an MS/MS resolution set to 17 500, a precursor automatic gain control target set to 5e4, a precursor isolation width set to 1.5 Da, and a maximum injection time set to 120 ms. The data was converted from .raw to .mgf file formats using ProteoWizard and analyzed through the Advanced Proteomics Facility Pipeline (CPFP; see (Trudgian et al., 2010)). A list of proteins was generated using CPFP meta-search Simple Protein Overview tool, using *L. mexicana* proteome as reference (gene models based on (Fiebig et al., 2015)) and label-free SINQ quantification, using search parameters outlined by (Beneke et al., 2019). Proteins were filtered based on a 5% false discovery rate (FDR) and the identification of at least 2 unique peptides.

### Immunoprecipitation

Cell lines expressing *LmxFused*::eYFP or *LmxULK4*::3xMYC were generated by tagging both alleles of the target gene with eYFP or 3xMYC, respectively, using the LeishGEdit method (Beneke et al., 2017). Specific primers were designed for amplification of donor DNA cassettes, as follows. To tag the C-terminus of *LmxULK4* with three MYC epitope tags, donor PCR products amplified from the standard pPLOT plasmids;

F 5’-GAGGCGAACAAGATTCTGAAGAAACTCTCCGGATCCGGATCAGGATCTGG-3’ R 5’-AAATCGTACAGGCAACAGCAAACCCGCCCGCCAATTTGAGAGACCTGTGC-3’.

To generate donor PCR products for *LmxFused* C-terminal tagging with an eYFP tag without a MYC epitope, eYFP was amplified from the pJ1170/pLEnT-YB plasmid (Dean et al., 2015):

F 5’-TACGTTGGCAACCTCGTCGATCGACTTGGTGGTTCTGGTAGTGGTTCCGGTTCCGGTTCTGTGAGCAAGGGCGAGGAGCTGTT-3’

R 5’-AAATCTGGAAAACGGCAACTATGAAGTCCGTTAGCCCTCCCACACATAACCAGAG-3’).

Immunoprecipitation with anti-GFP antibodies followed the protocol from Akiyoshi & Gull (2014), with the following modifications: Instead of using PEME buffer, cells were extracted directly in modified buffer H (BH)/0.15 (25 mM HEPES pH 8.0, 2 mM MgCl_2_, 0.1 mM EDTA pH 8.0, 0.5 mM EGTA pH 8.0, 1% NP-40, 150 mM KCl, and 15% glycerol), including protease inhibitors (Leupeptin, Pepstatin, E-64, 20 μg/ml each, and 0.2 mM PMSF) and phosphatase inhibitors (1 mM sodium pyrophosphate, 2 mM Na-beta-glycerophosphate, 0.1 mM Na_3_VO_4_, 5 mM NaF, and 100 nM microcystin-LR). Cell extracts were frozen in liquid N_2_ and ground in a mortar and pestle, without sonication. Cell extracts were then centrifuged (14000*g* for 30 minutes at 4°C) and only the supernatant was forwarded to immunoprecipitation. Post-IP bead samples were then washed twice in PBS and eluted in 0.2M Glycine, pH 2.5 (shaking at 21°C for 7 minutes) and neutralised with 10% volume 1M Tris-HCl, pH 8.8. Equivalent proportions of each sample were collected at multiple points during the protocol, with SDS-PAGE and immunoblots performed by standard methods using the following mouse monoclonal antibodies: anti-GFP (11814460001, Roche) and anti-MYC (clone 4A6, 05-724, Merck).

### LC-MS/MS for IP/Immunoprecipitation

Mass spectrometry analysis was performed by the Advanced Proteomics facility (Department of Biochemistry, University of Oxford). Peptides were separated by nano liquid chromatography (Thermo Scientific Ultimate RSLC 3000) coupled in line a Q Exactive mass spectrometer equipped with an Easy-Spray source (Thermo Fischer Scientific). Peptides were trapped onto a C18 PepMac100 precolumn (300µm i.d.x5mm, 100Å, ThermoFischer Scientific) using Solvent A (0.1% Formic acid, HPLC grade water). The peptides were further separated onto an Easy-Spray RSLC C18 column (75um i.d., 50cm length, Thermo Fischer Scientific) using a 60 minutes linear gradient (15% to 35% solvent B (0.1% formic acid in acetonitrile)) at a flow rate 200nl/min. The raw data were acquired on the mass spectrometer in a data-dependent acquisition mode (DDA). Full-scan MS spectra were acquired in the Orbitrap (Scan range 350-1500m/z, resolution 70,000; AGC target, 3e6, maximum injection time, 50ms). The 10 most intense peaks were selected for higher-energy collision dissociation (HCD) fragmentation at 30% of normalized collision energy. HCD spectra were acquired in the Orbitrap at resolution 17,500, AGC target 5e4, maximum injection time 120ms with fixed mass at 180m/z. Charge exclusion was selected for unassigned and 1+ ions. The dynamic exclusion was set to 20 s.

## Acknowledgements

We thank Sabrina Liberatori and Marjorie Fournier (University of Oxford Proteomics facility) for mass spectrometry support, Raman Dhaliwal and Errin Johnson (Dunn School Bioimaging facility) for electron microscopy support, James Smith (University of Oxford) for help with the generation of *Leishmania* mutants, Richard Wheeler (University of Oxford) for advice on IFT measurements, Bungo Akiyoshi (University of Oxford) for advice on the IP protocol, Alice Meunier and Nathalie Spassky (ENS Paris) for introducing us to preparation of ependymal cell cultures, Jeremy Mottram and Vincent Geoghegan (University of York) for sharing their XL-BioID protocol prior to publication and Keith Gull (University of Oxford) for antibody C3B9, access to laboratory equipment and helpful comments on the manuscript.

## Competing Interests

No competing interests declared

## Funding

CMC and EG were funded by the UK Medical Research Council (MRC) and the UK Department for International Development (DFID) under the MRC/DFID Concordat agreement; grant no. MR/R000859/1 (https://mrc.ukri.org/).

EG was supported by a Royal Society University Research Fellowship (UF160661; https://royalsociety.org/)

HPV was supported by an ERASMUS+ mobility grant

TB was supported by MRC PhD studentship (15/16_MSD_836338)

VV laboratory was supported by the Czech Science Foundation (GA CR) project no. 20-23165J and by an Installation Grant from the European Molecular Biology Organization.

PG is a student of the Faculty of Science, Charles University, Prague, which provided a PhD student fellowship.

We acknowledge support for this project for EG through the WCIP core Wellcome Centre Award [104111/Z/14/Z] and through the Wellcome Trust grant [104627/Z/14/Z] to Keith Gull; https://wellcome.org).

We acknowledge the Light Microscopy Core Facility, IMG CAS, Prague, Czech Republic, supported by MEYS (LM2018129, CZ.02.1.01/0.0/0.0/18_046/0016045) and RVO: 68378050-KAV-NPUI, for their support with the confocal imaging of ependymal cells.

The funders had no role in study design, data collection and analysis, decision to publish, or preparation of the manuscript.

## Notes

### Competing Interest Statement

The authors have declared no competing interest.

