## Supplemental Figures for "ULK4 and Fused/STK36 interact to mediate assembly of a motile flagellum"

McCoy *et al.* 2022

#### *Supplemental Data File 1*

Protein identification for BioID experiment (.xlsx file).

#### *Supplemental Data File 2*

Protein identification for Immunoprecipitation experiment (.xlsx file).

#### *Supplemental Data File 3*

Primer sequences used for validation of Gene KO by PCR (.xlsx file).

### Supplemental Figures

#### *S1 Fig. Loss of LmxFused::mNG and LmxULK4::mNG signal from detergent-extracted cells.*

**(A)** Top panel, live promastigotes expressing LmxULK4::mNG, imaged by phase contrast and fluorescence microscopy. Bottom panel, the same cell line, imaged after treatment with 0.1% IGEPAL® CA-630. **(B)** Top panel, live promastigotes expressing LmxFused::mNG. Bottom panel, the same cell line, after treatment with 0.1% IGEPAL® CA-630. Hoechst 33342 staining indicates nuclear and kinetoplast DNA.

#### *S2 Fig. Verification of KO cell lines.*

(**A**) Cartoons depicting diagnostic PCR strategy to confirm gene KO though amplification of a fragment of the target gene ORF. (**B**) Products from target gene ORF PCR, run on an agarose gel. Each panel shows the product obtained from the putative knockout cell line (KO) and the parental cell line (P). Empty lanes are marked with "**-**". 'Gene target' indicates name or GeneID of the KO target gene. 'Parental' indicates cell line in which the gene was targeted, see main text for details of cell lines. An asterisk replaces the 'LmxM.' GeneID prefix. Absence of the target ORF PCR product was confirmed for all genes. See S3 Table for primers used. Fainter bands below the target gene amplicon are likely primer dimers. (**C**) Cartoons depicting diagnostic PCR strategy to verify DNA sample quality through amplification of a fragment of the inserted blasticidin resistance gene (BlastR). (**D**) Products from BlastR PCR run on an agarose gel. (**E-F**) Reads from Illumina sequencing of genomic DNA from *ΔLmxFused* and

*ΔLmxULK4* cell lines, respectively, were mapped to the *L. mexicana* reference genome. The screenshots show the genomic locus for *LmxFused* (E) and *ΔLmxULK4* (F), visualised with the Integrative Genomics Viewer. *ΔLmxFused* indicates reads derived from the *ΔLmxFused* cell line, *ΔLmxULK4* indicates reads derived from the *ΔLmxULK4* cell line.

#### *S3 Fig. Growth curves of ΔLmxULK4 and ΔLmxFused cell lines.*

**(A-B)** Δ*LmxULK4* and Δ*LmxFused* promastigotes and the parental *L. mex* Cas9 T7 line were grown in M199 medium and diluted into fresh medium every 24 hours to allow for sustained exponential growth. (**C**) Parasite doubling times, calculated from the data in (**A-B**). (**D**) Growth curve of Δ*LmxULK4,* Δ*LmxFused* and *L. mex* Cas9 T7 promastigotes in continuous culture.

#### *S4 Fig. Persistence of LmxULK4::mNG and LmxFused::mNG signal in ΔPF20 cell lines.*

Visualisation of LmxULK4::mNG (**A**) and LmxFused::mNG (**B**) in live promastigotes before and after deletion of *PF20*. The composite images show overlays of phase contrast images with the green fluorescence channels and fluorescence of Hoechst 33342 to indicate nuclear and kinetoplast DNA.

#### *S5 Fig. Structural defects at the base of mutant flagella.*

**(A)** Parental control. TEM serial sections through the base of the flagellum, showing microtubule triplets of the basal body (1), doublet microtubules in the transition zone (4-5) and the symmetrical 9+2 microtubule axoneme emerging into the flagellar pocket (FP; 6-8). **(B,C)** Examples of Δ*LmxFused* cells with axoneme defects. In (B), a triplet microtubule basal body showing nine-fold symmetry is formed (3). Microtubules terminate at the transition zone (5) with only one or two microtubules extending slightly further (6; white arrowheads) before ending, still within the FP. Membrane-bounded structures of unknown origin are observed in the FP (7,8 asterisks). In (C) only four doublet microtubules extended beyond the transition zone (6), of which only three extended to the distal-most section captured in this series (8). **(D)** Example of cell where axoneme and central pair microtubules are not formed correctly. This cell has a basal body with only eight triplet microtubules (3), extending though a transition zone (4-5) and axoneme with eight doublet microtubules (6-8; black arrowheads), with only one singlet microtubule in the centre that originates from the basal plate is seen in (5). **(E)** Example of a cell where the nine-fold radial symmetry of the axoneme is lost. Doublet microtubules and two singlets are seen extending from the basal plate (1) though the FP (2-8). Eight doublets appear normal in cross-section (black arrowheads in 4), the ninth doublet (white arrowheads) appears to be present at the basal plate but is either mispositioned such that it cannot be seen clearly, or structurally incomplete.

**_S6 Fig. Measurement of IFT in ΔLmxULK4 and ΔLmxFused cell lines._**

(**A-C**) Frequency distribution of IFT particle velocity in the parental cell line *L. mex* Cas9 T7 mNG::IFT81 (A), and the derived KO lines Δ*LmxULK4* (B) and Δ*LmxFused* (C). **(D-F)** Particle intensity plotted against velocity for the mNG::IFT81 parental line (D), Δ*LmxULK4* (E) and Δ*LmxFused* (F). **(G)** Summary of the mNG::IFT81 particle replicate number and mean velocities recorded. **(H-J)** Frequency distribution of IFT particle velocity in the parental cell line *L. mex* Cas9 T7 mNG::IFT140 (H), and the derived KO cell lines Δ*LmxULK4* (I) and Δ*LmxFused* (J). **(K-M)** Particle intensity plotted against velocity for the mNG::IFT140 parental line (K), Δ*LmxULK4* (I) and Δ*LmxFused* (M). **(N)** Summary of the mNG::IFT140 particle replicate number and mean velocities recorded. **(O)** Localisation of mNG::IFT81. **(P)** Localisation of mNG::IFT140. Panels in (O-P) show the localisation of the respective IFT protein in the parental cell line and one example cell from each of the following distinct categories observed in Δ*LmxULK4* and Δ*LmxFused* cell lines: (i) IFT signal within a long external flagellum extending outside of the flagellar pocket, (ii) IFT signal running along the length of the flagellum when restricted to the flagellar pocket, (iii) an enriched mNG spot localised near the kinetoplast DNA (assumed to be at the basal body) or (iv) an apparent lack of enrichment in flagellum associated IFT signal. Red arrows indicate the absence of IFT signal at the basal body region. The percentage of cells in each category is indicated ($n \geq 500$). Nuclear and kinetoplast DNA are labelled with Hoechst 33342.

**_S7 Fig. Characterisation of the LmxFused::BirA* cell line._**

**(A)** Growth of LmxFused::BirA*::MYC expressing promastigotes in M199 medium. **(B)** Immuno-fluorescence detection of LmxFused::BirA* in promastigotes, using anti-MYC (4A6) antibody against the fusion protein and a TRITC-conjugated secondary antibody. Red arrows point to the tip of the flagellum. **(C)** Detection of LmxFused::BirA*::MYC on a western blot of whole cell protein lysates (‘before beads’) and samples taken at different steps during the streptavidin bead mediated pulldown. Numbers indicate molecular weight in kDa.

**_S8 Fig. Characterisation of the LmxFused::eYFP cell line._**

**(A)** Localisation of LmxFused::eYFP in *Leishmania* promastigotes. Red arrows highlight the base and tip of the flagellum of a non-dividing 1K1N1F cell that display eYFP signal enrichment. Nuclear and kinetoplast DNA were labelled with Hoechst 33342. **(B)** Western blot of proteins samples from the parental cell line and from cells expressing LmxFused::eYFP, taken at different steps of the immuno-precipitation procedure and probed with anti-GFP antibody. Numbers indicate molecular weight (MW) in kDa; the calculated MW for

LmxFused::eYFP is 147 kDa. The red arrow points to an additional lower molecular weight band detected with the antibody, which may represent a fragment of LmxFused::eYFP. **(C)** Western blot of proteins samples from the parental cell line and from cells expressing LmxULK4::MYC, taken at different steps of the immuno-precipitation procedure and probed with anti-MYC antibody. The black arrow points to a band of the expected MW for LmxULK4::MYC (116 kDa). Additional observed bands suggest cleavage or degradation of LmxULK4::MYC occurred, following cell lysis. Red arrows point to lower molecular weight MYC epitope positive bands that are seen in the cell lysate but were not detected in the whole cell sample. **(D)** Western blot showing pulldown and elution of the GFP control bait protein. **(E)** LmxULK4 peptides identified via mass spectrometry in the LmxFused::eYFP sample, aligned to the whole LmxULK4 protein.

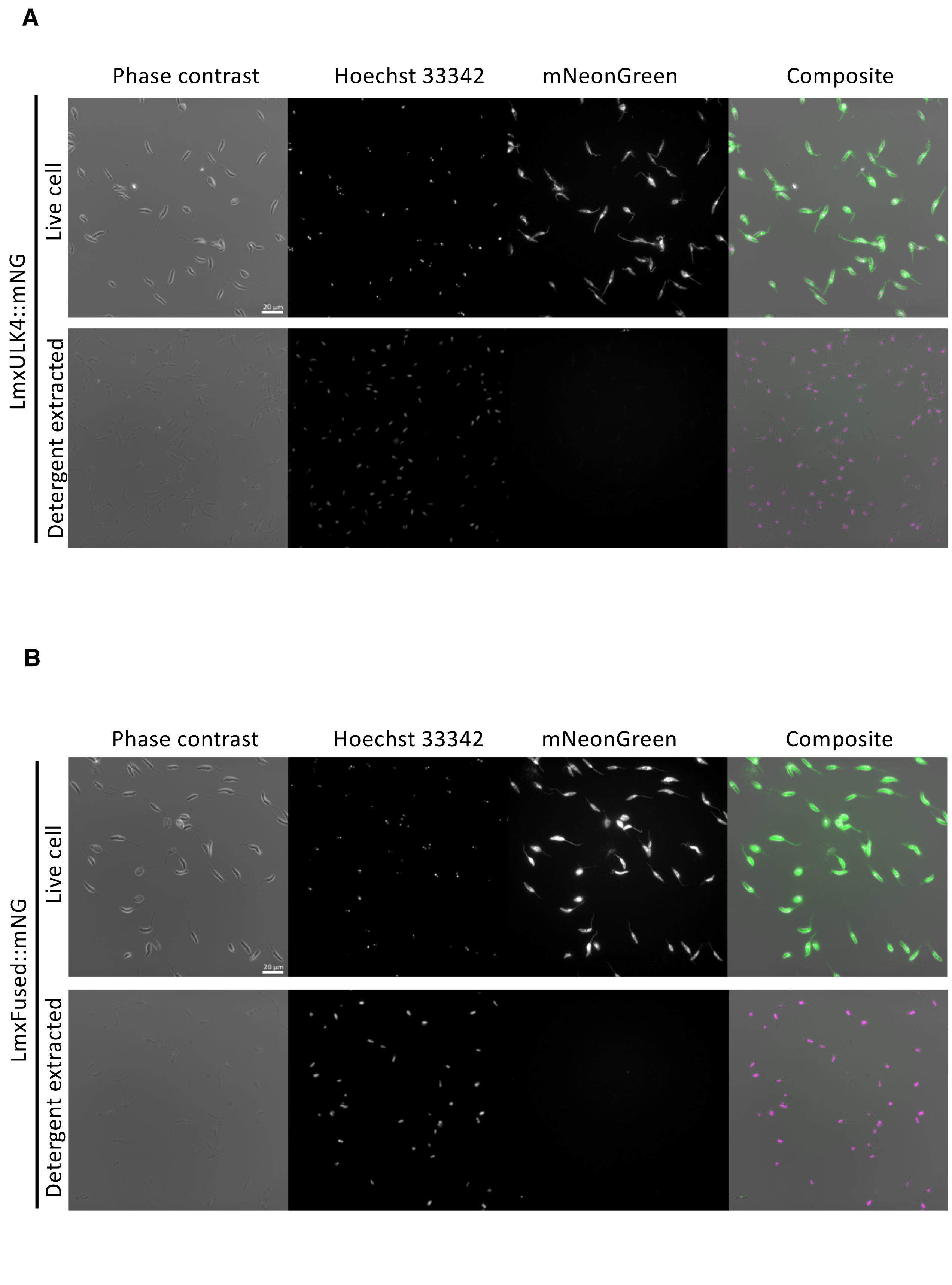
A
Phase contrast
Hoechst 33342
mNeonGreen
Composite
LmxULK4::mNG
Live cell
Detergent extracted
20 µm
B
Phase contrast
Hoechst 33342
mNeonGreen
Composite
LmxFused::mNG
Live cell
Detergent extracted
20 µm

S Figure 1

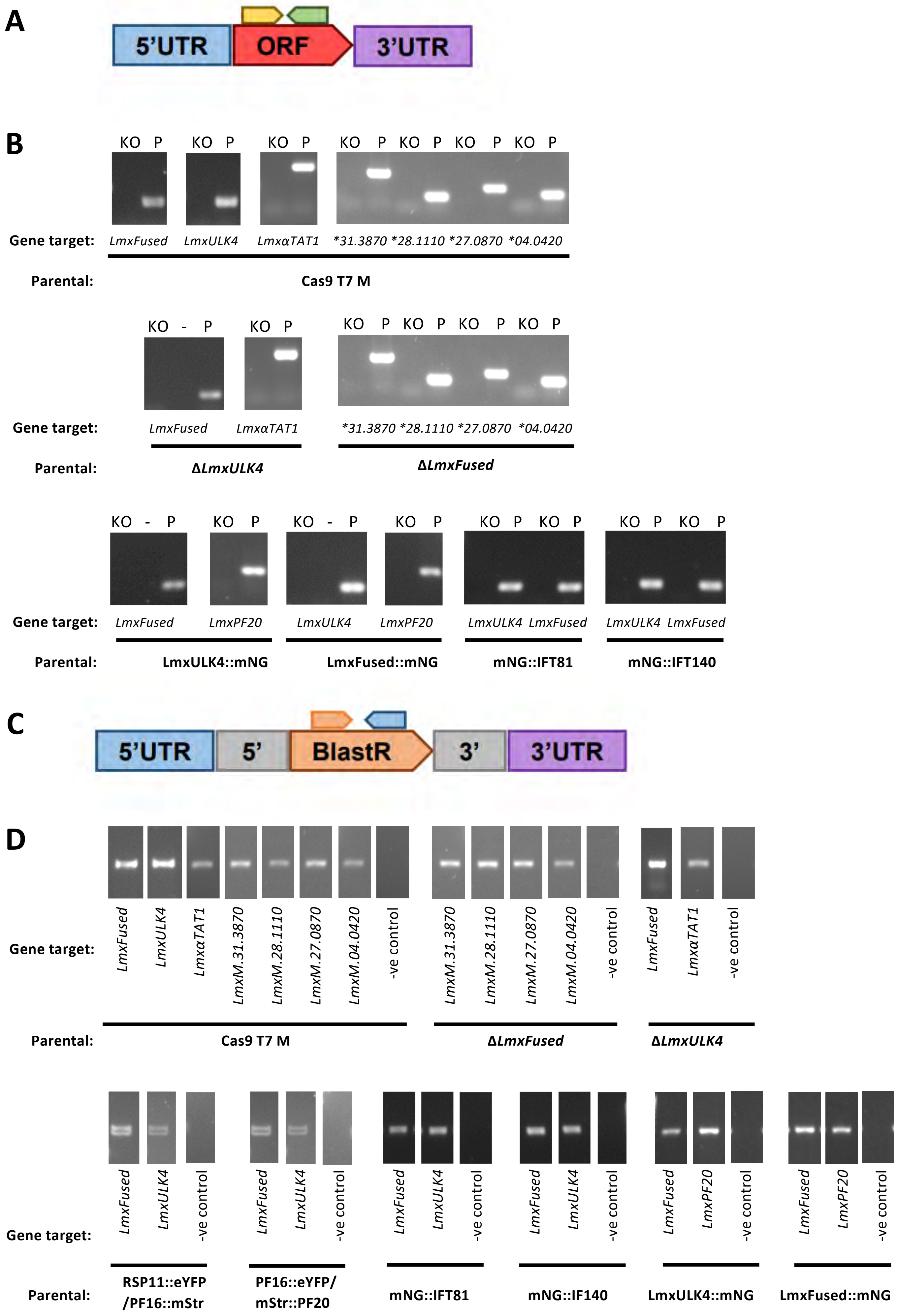
A
5'UTR
ORF
3'UTR
B
KO P KO P KO P KO P KO P KO P KO P
Gene target:
LmxFused LmxULK4 LmxαTAT1 *31.3870 *28.1110 *27.0870 *04.0420
Parental:
Cas9 T7 M
KO - P KO P KO P KO P KO P KO P
Gene target:
LmxFused LmxαTAT1 *31.3870 *28.1110 *27.0870 *04.0420
Parental:
ΔLmxULK4
ΔLmxFused
KO - P KO P KO - P KO P KO P KO P KO P KO P
Gene target:
LmxFused LmxPF20 LmxULK4 LmxPF20 LmxULK4 LmxFused LmxULK4 LmxFused
Parental:
LmxULK4::mNG
LmxFused::mNG
mNG::IFT81
mNG::IFT140
C
5'UTR
5'
BlastR
3'
3'UTR
D
Gene target:
LmxFused LmxULK4 LmxαTAT1 LmxM.31.3870 LmxM.28.1110 LmxM.27.0870 LmxM.04.0420 -ve control
LmxM.31.3870 LmxM.28.1110 LmxM.27.0870 LmxM.04.0420 -ve control
LmxFused LmxαTAT1 -ve control
Parental:
Cas9 T7 M
ΔLmxFused
ΔLmxULK4
Gene target:
LmxFused LmxULK4 -ve control
LmxFused LmxULK4 -ve control
LmxFused LmxULK4 -ve control
LmxFused LmxULK4 -ve control
LmxFused LmxPF20 -ve control
LmxFused LmxPF20 -ve control
Parental:
RSP11::eYFP /PF16::mStr
PF16::eYFP/ mStr::PF20
mNG::IFT81
mNG::IF140
LmxULK4::mNG
LmxFused::mNG

S Figure 2

E

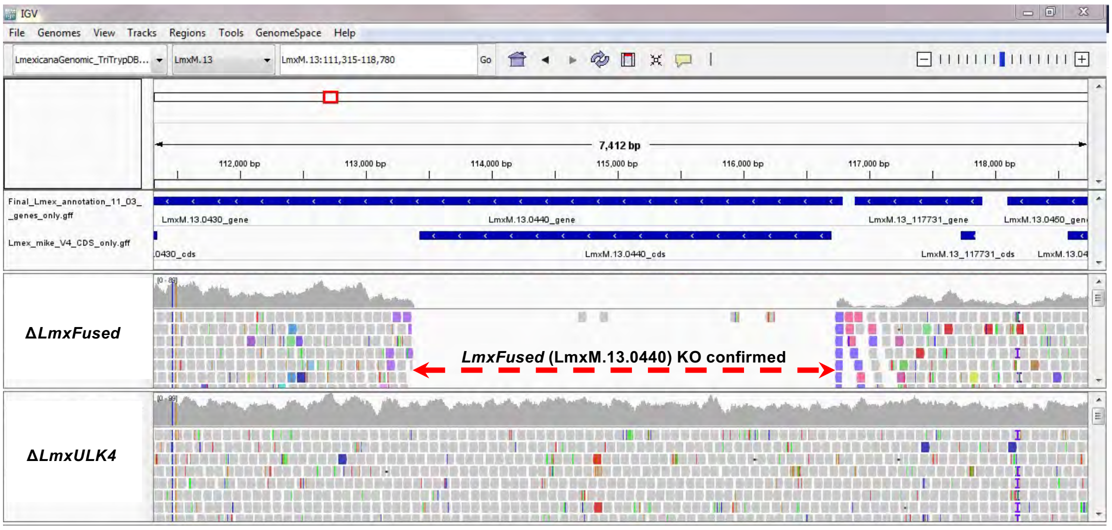
IGV
File Genomes View Tracks Regions Tools GenomeSpace Help
LmexicanaGenomic_TriTrypDB...
LmxM.13
LmxM.13:111,315-118,780
Go
7,412 bp
112,000 bp
113,000 bp
114,000 bp
115,000 bp
116,000 bp
117,000 bp
118,000 bp
Final_Lmex_annotation_11_03_
_genes_only.gff
Lmex_mike_V4_CDS_only.gff
LmxM.13.0430_gene
LmxM.13.0440_gene
LmxM.13_117731_gene
LmxM.13.0450_gene
.0430_cds
LmxM.13.0440_cds
LmxM.13_117731_cds
ΔLmxFused
LmxFused (LmxM.13.0440) KO confirmed
ΔLmxULK4

F

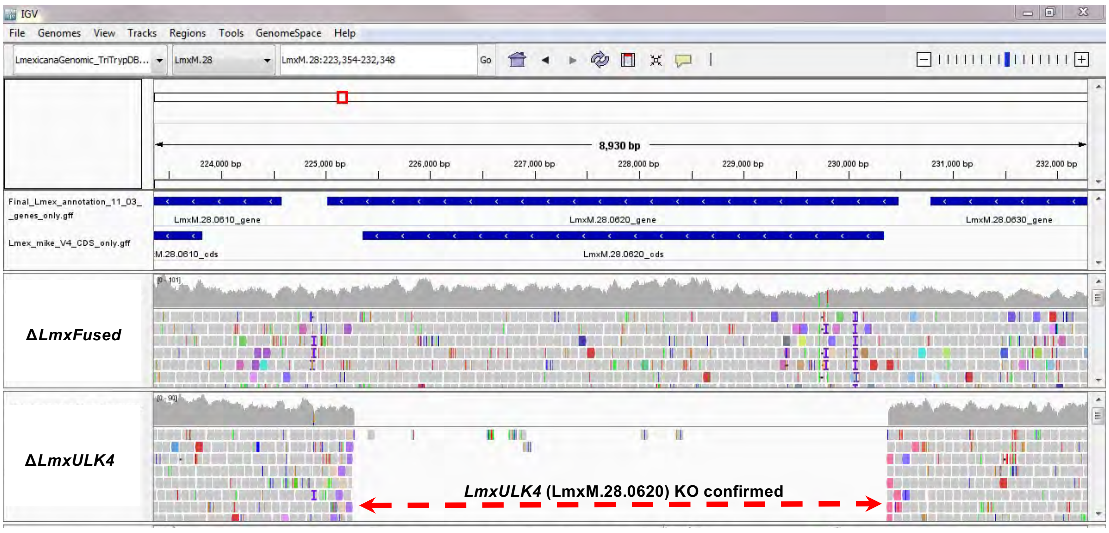
IGV
File Genomes View Tracks Regions Tools GenomeSpace Help
LmexicanaGenomic_TriTrypDB...
LmxM.28
LmxM.28:223,354-232,348
Go
8,930 bp
224,000 bp
225,000 bp
226,000 bp
227,000 bp
228,000 bp
229,000 bp
230,000 bp
231,000 bp
232,000 bp
Final_Lmex_annotation_11_03_
_genes_only.gff
Lmex_mike_V4_CDS_only.gff
LmxM.28.0610_gene
LmxM.28.0620_gene
LmxM.28.0630_gene
M.28.0610_cds
LmxM.28.0620_cds
ΔLmxFused
ΔLmxULK4
LmxULK4 (LmxM.28.0620) KO confirmed

S Figure 2

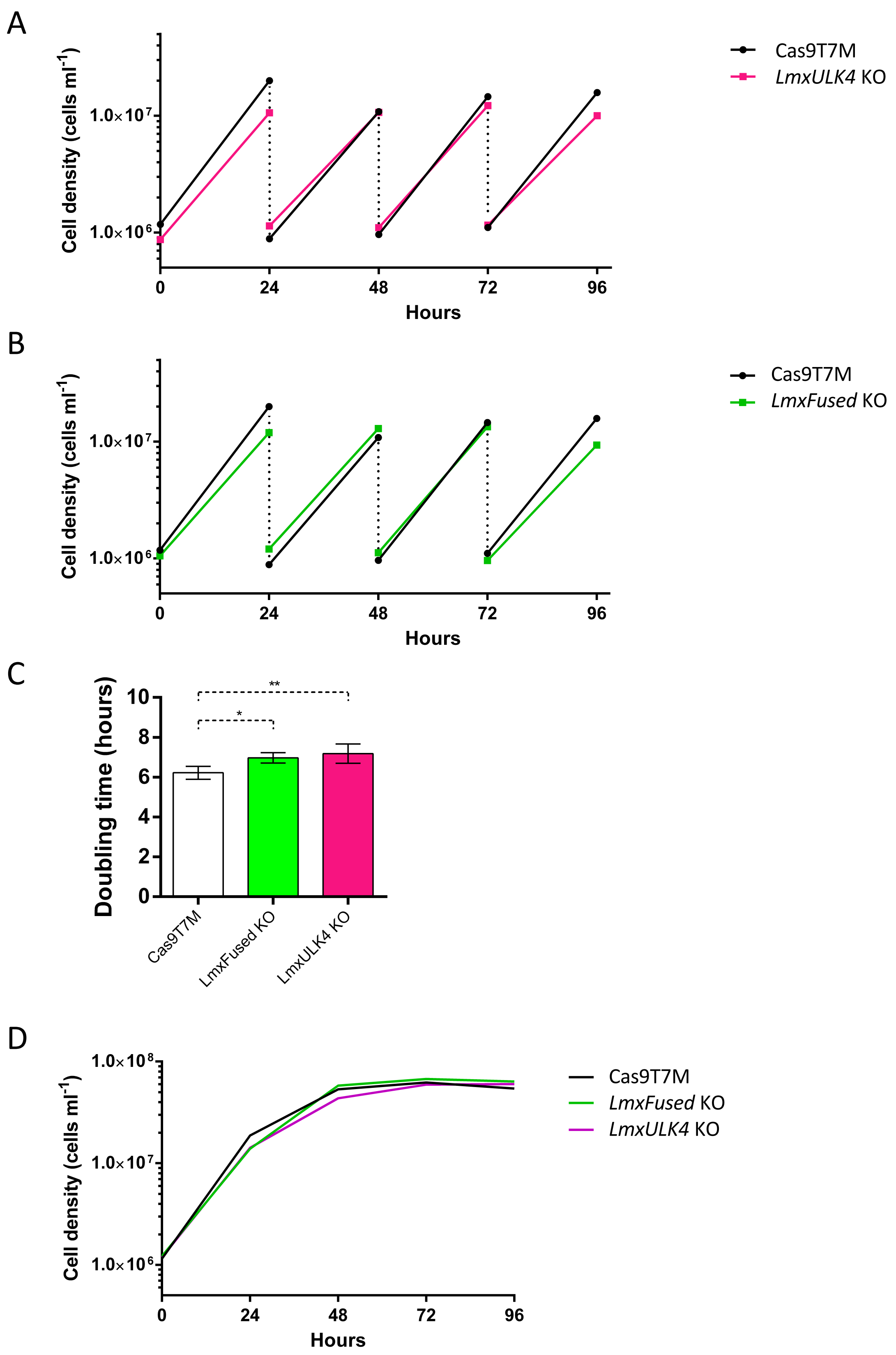
A
Cell density (cells ml-1)
1.0×10^7
1.0×10^6
0
24
48
72
96
Hours
Cas9T7M
LmxULK4 KO
B
Cell density (cells ml-1)
1.0×10^7
1.0×10^6
0
24
48
72
96
Hours
Cas9T7M
LmxFused KO
C
Doubling time (hours)
10
8
6
4
2
0
**
*
Cas9T7M
LmxFused KO
LmxULK4 KO
D
Cell density (cells ml-1)
1.0×10^8
1.0×10^7
1.0×10^6
0
24
48
72
96
Hours
Cas9T7M
LmxFused KO
LmxULK4 KO

S Figure 3

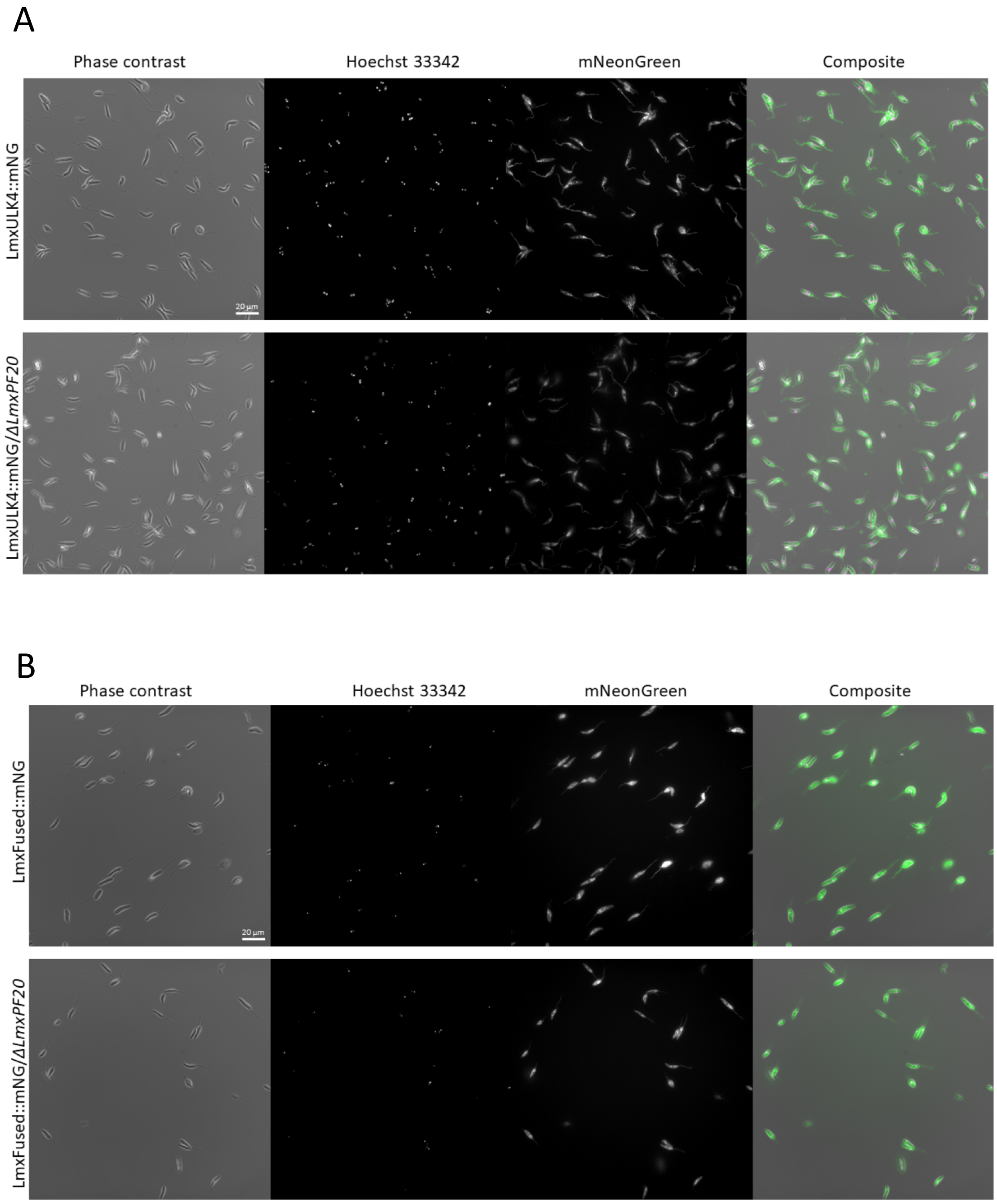
A
Phase contrast
Hoechst 33342
mNeonGreen
Composite
LmxULK4::mNG
LmxULK4::mNG/ΔLmxPF20
20 µm
B
Phase contrast
Hoechst 33342
mNeonGreen
Composite
LmxFused::mNG
LmxFused::mNG/ΔLmxPF20
20 µm

S Figure 4

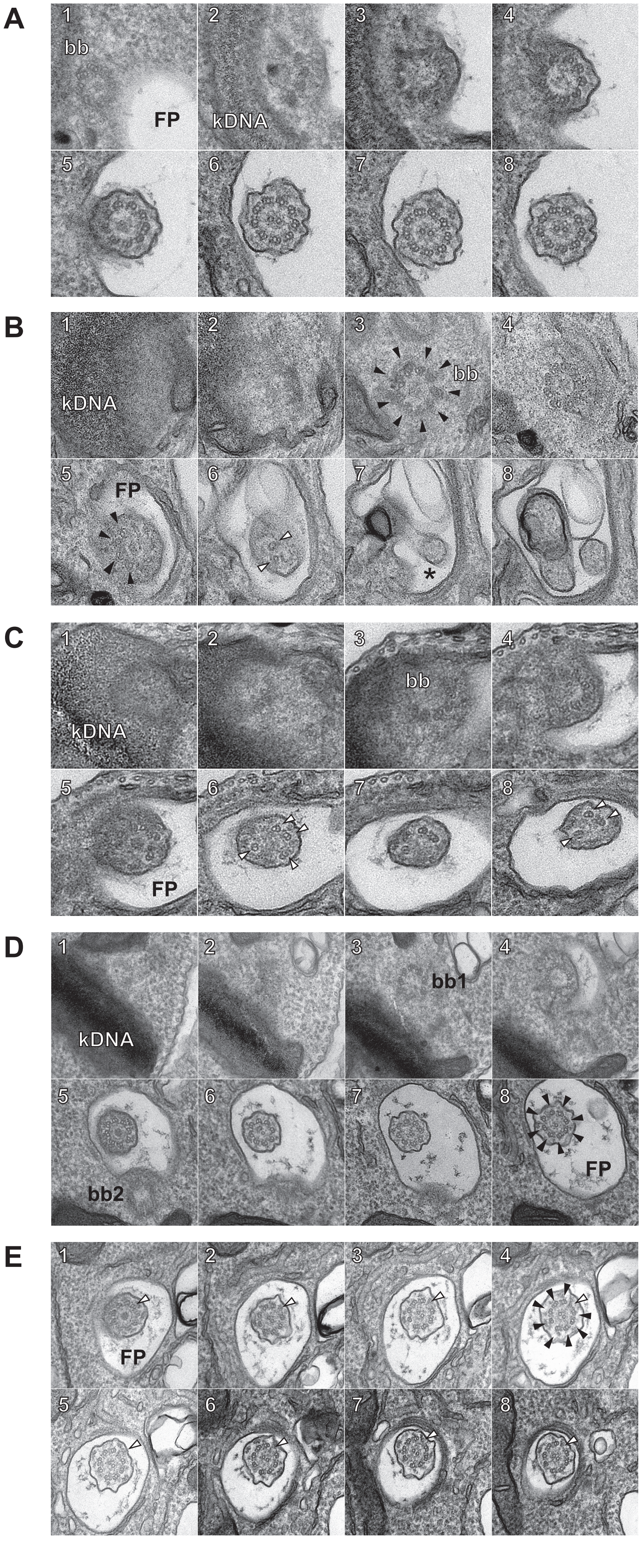

A
1
bb
FP
2
kDNA
3
4
5
6
7
8
B
1
kDNA
2
3
bb
4
5
FP
6
7
*
8
C
1
kDNA
2
3
bb
4
5
FP
6
7
8
D
1
kDNA
2
3
bb1
4
5
bb2
6
7
8
FP
E
1
FP
2
3
4
5
6
7
8


S Figure 5

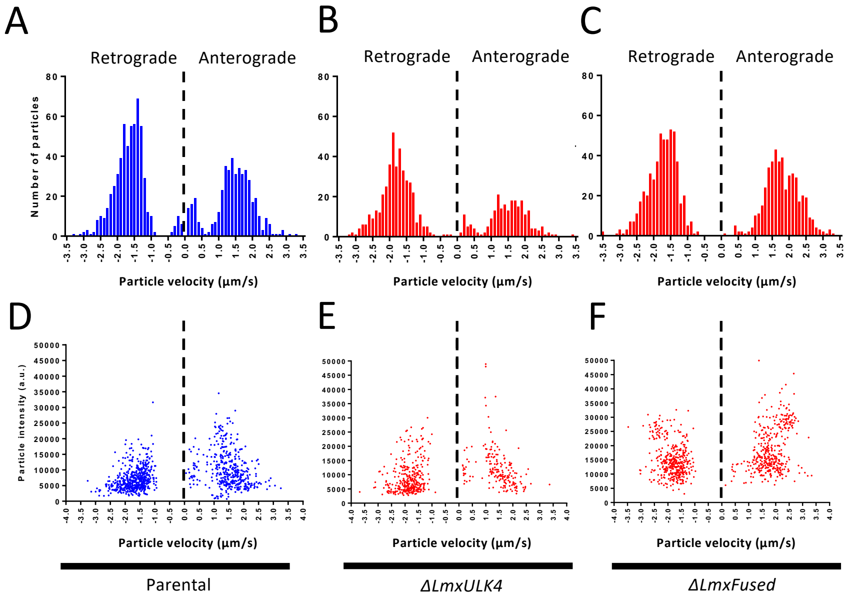


G

| Cell lines | Anterograde particle number | Retrograde particle number | Cell number | Anterograde velocity (µm/s) | Retrograde velocity (µm/s) |
|---|---|---|---|---|---|
| Parental | 447 | 579 | 16 | 1.42 ± 0.61 | -1.63 ± 0.48 |
| *ΔLmxULK4* | 239 | 426 | 15 | 1.46 ± 0.61 | -1.84 ± 0.50 |
| *ΔLmxFused* | 467 | 531 | 14 | 1.80 ± 0.51 | -1.76 ± 0.45 |

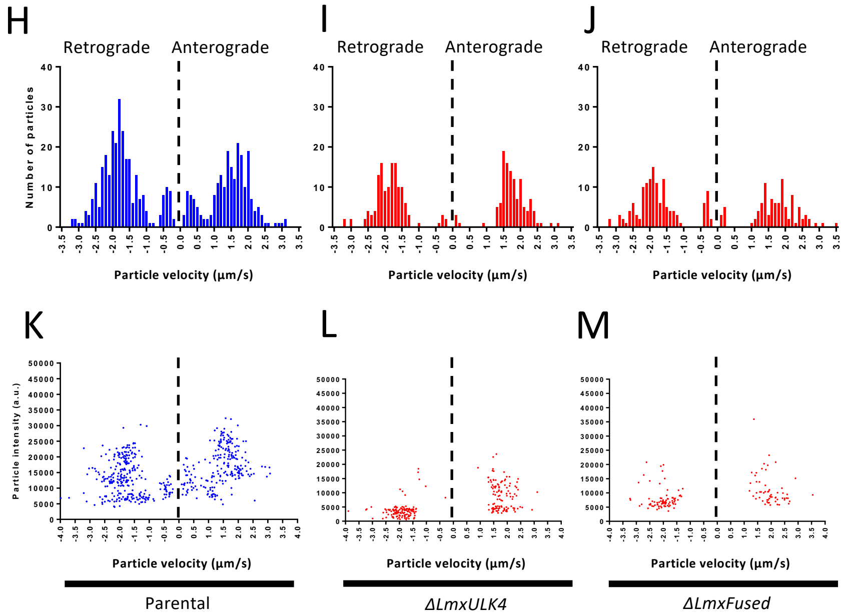


N

| Cell lines | Anterograde particle number | Retrograde particle number | Cell number | Anterograde velocity (µm/s) | Retrograde velocity (µm/s) |
|---|---|---|---|---|---|
| Parental | 224 | 291 | 13 | 1.44 ± 0.63 | -1.73 ± 0.65 |
| *ΔLmxULK4* | 120 | 134 | 13 | 1.74 ± 0.46 | -1.86 ± 0.54 |
| *ΔLmxFused* | 99 | 134 | 13 | 1.76 ± 0.75 | -1.76 ± 0.69 |

S Figure 6

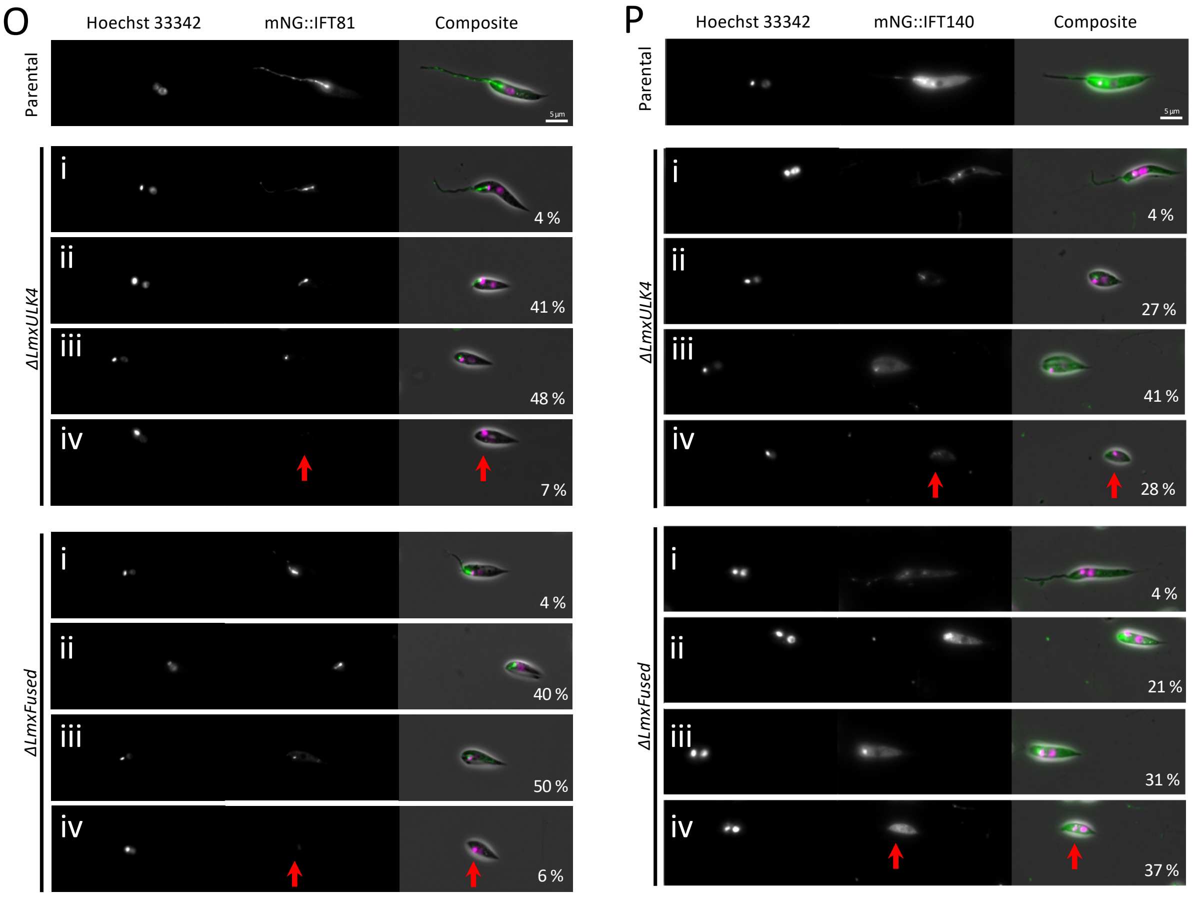
O
Hoechst 33342
mNG::IFT81
Composite
Parental
ΔLmxULK4
i
4 %
ii
41 %
iii
48 %
iv
7 %
ΔLmxFused
i
4 %
ii
40 %
iii
50 %
iv
6 %
P
Hoechst 33342
mNG::IFT140
Composite
Parental
ΔLmxULK4
i
4 %
ii
27 %
iii
41 %
iv
28 %
ΔLmxFused
i
4 %
ii
21 %
iii
31 %
iv
37 %

S Figure 6

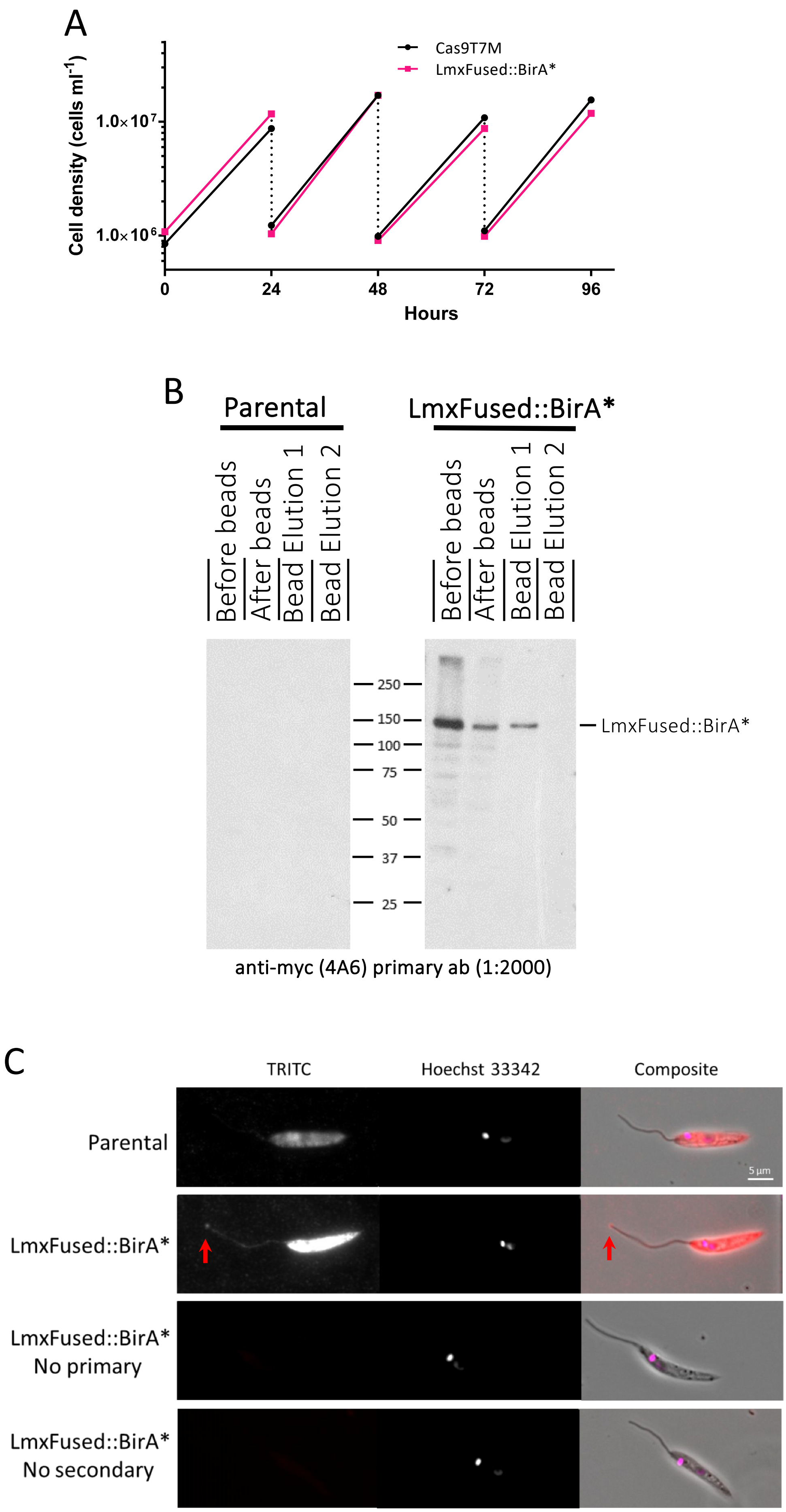
A
Cas9T7M
LmxFused::BirA*
Cell density (cells ml-1)
1.0×10^7
1.0×10^6
0
24
48
72
96
Hours
B
Parental
LmxFused::BirA*
Before beads
After beads
Bead Elution 1
Bead Elution 2
Before beads
After beads
Bead Elution 1
Bead Elution 2
250
150
100
75
50
37
25
LmxFused::BirA*
anti-myc (4A6) primary ab (1:2000)
C
TRITC
Hoechst 33342
Composite
Parental
5 µm
LmxFused::BirA*
LmxFused::BirA*
No primary
LmxFused::BirA*
No secondary

S Figure 7

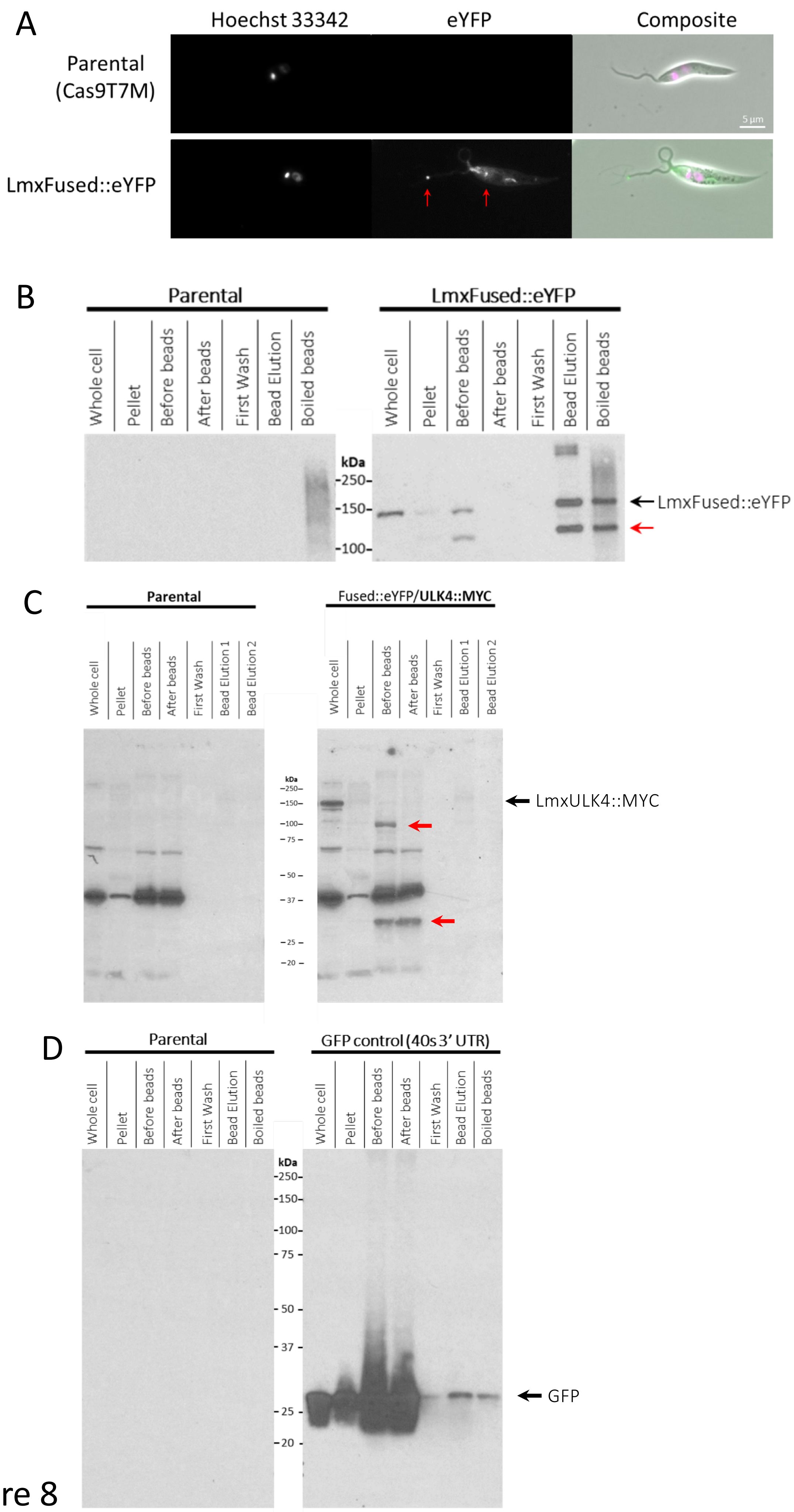
A
Hoechst 33342
eYFP
Composite
Parental (Cas9T7M)
LmxFused::eYFP
5 µm
B
Parental
LmxFused::eYFP
Whole cell
Pellet
Before beads
After beads
First Wash
Bead Elution
Boiled beads
kDa
250
150
100
LmxFused::eYFP
C
Parental
Fused::eYFP/ULK4::MYC
Whole cell
Pellet
Before beads
After beads
First Wash
Bead Elution 1
Bead Elution 2
LmxULK4::MYC
D
Parental
GFP control (40s 3' UTR)
Whole cell
Pellet
Before beads
After beads
First Wash
Bead Elution
Boiled beads
kDa
250
150
100
75
50
37
25
20
GFP

S Figure 8

E

LmxULK4 N-terminus

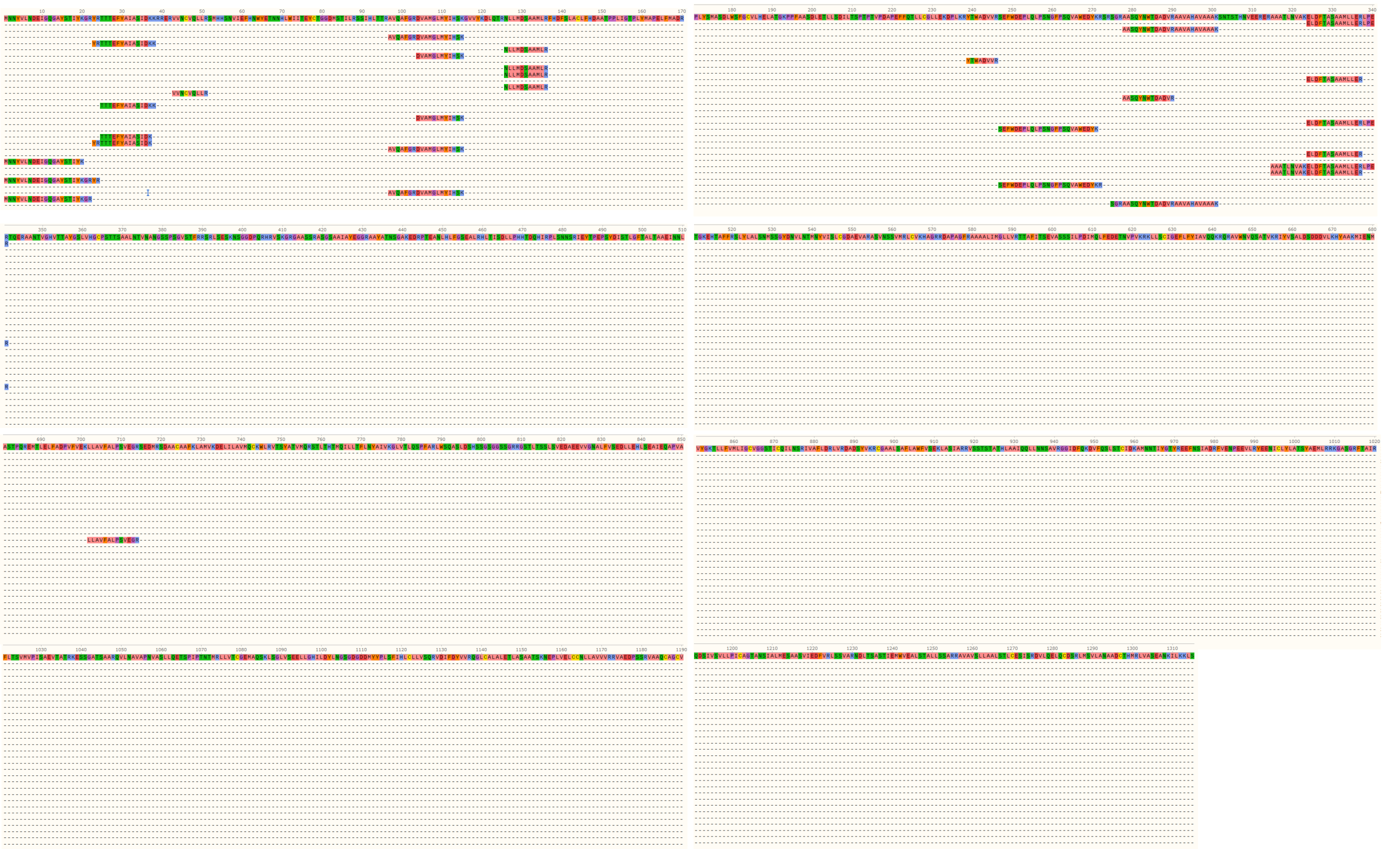

C-terminus

S Figure 8
